# “Rapid prototyping of flexible biodegradable ECoG arrays for high-resolution cortical mapping and real-time seizure classification ”

**DOI:** 10.1101/2025.11.02.686028

**Authors:** Sahera Saleh, Nisrine Bakri, Khouloud Issa, Heba Badawe, Tamara Al-Sadek, Reem El Jammal, Mariette Awad, Makram Obeid, Massoud Khraiche

## Abstract

Neural interfaces are essential tools for diagnosing and managing neurological disorders, yet conventional electrocorticography (ECoG) devices are limited by mechanical mismatch with brain tissue, chronic inflammation, and poor scalability. Here, we introduce a fully inkjet-printed, flexible, and biodegradable ECoG array fabricated on ultrathin polycaprolactone films with gold nanoparticle electrodes. The arrays achieve among the highest electrode densities reported for additive manufacturing (7.44 electrodes/mm²) while maintaining low impedance (10.6 kΩ at 1 kHz) and high-fidelity recordings (SNR 28 dB). A rapid, maskless prototyping process relying on photonic sintering enables scalable, cost-effective fabrication. In vivo, the devices conformally mapped cortical seizure propagation and resolved distinct ictal dynamics in a rat model. Histology at 30 days confirmed preserved neuronal density and astrocytic response comparable to controls, indicating minimal chronic inflammation. A convolutional neural network trained on recorded signals classified seizure stages with >95% accuracy, underscoring the translational potential for real-time monitoring and closed-loop neuromodulation. This platform unites rapid prototyping, biodegradability, and high performance, providing a scalable route toward next-generation, patient-specific, and disposable neural interfaces for epilepsy and other neuroengineering applications.

## Introduction

Neural interface technologies play a central role in understanding brain function and treating neurological disorders. By recording or stimulating neural activity, they provide insights into disease mechanisms, guide surgical interventions, and enable neuroprosthetic control. Among available modalities, electroencephalography (EEG) and penetrating microelectrode arrays (MEAs) represent two ends of the spectrum [1]. EEG offers non-invasive coverage of large cortical areas but suffers from poor spatial resolution due to signal attenuation by scalp and skull [2–4]. MEAs, by contrast, access individual neurons but their invasiveness often leads to gliosis, tissue damage, and long-term instability [5–7].

Electrocorticography (ECoG) serve as an effective middle ground, offering higher signal fidelity than EEG with reduced invasiveness compared to MEAs. ECoG arrays are used clinically for pre-surgical epilepsy monitoring, functional mapping, and increasingly for brain–computer interfaces (BCIs) [8–11]. Despite these advances, current devices rely on rigid metallic electrodes embedded in silicone or polyimide substrates. The mechanical mismatch between such materials and the soft, dynamic brain surface leads to micromotion-induced artifacts, chronic inflammation, and reduced stability over time [12, 13]. Furthermore, conventional devices are not designed for resorption or disposal, making explantation necessary once their clinical role ends. Their fabrication, typically reliant on cleanroom lithography, is costly and inflexible, hindering rapid adaptation for patient-specific applications.

Recent advancements in flexible bioelectronics have enabled the development of next-generation ECoG arrays with improved mechanical conformability. Ultra-thin, stretchable electrodes fabricated from biocompatible polymers (e.g., parylene-C, polyimide, silk fibroin) and emerging soft materials such as graphene and organic conductors have demonstrated enhanced biostability and recording fidelity [8, 14–16]. Despite these advances, traditional microfabrication techniques remain complex, time-consuming, and costly, limiting scalability for broader clinical and research applications. Moreover, implantable ECoG arrays often require surgical retraction once seizure localization is complete or due to long-term biocompatibility concerns. This necessity highlights the need for bioresorbable or easily retrievable neural interfaces to minimize surgical risks and patient burden [16–18].

Additive manufacturing techniques such as inkjet printing have been explored as scalable and cost-effective approaches for fabricating neural interfaces. Inkjet printing enables precise, maskless deposition of conductive, dielectric and insulating materials onto flexible substrates, simplifying fabrication and reducing material waste [2, 19, 20]. While this approach facilitates rapid prototyping and customization of ECoG arrays, many printed neural interfaces have yet to achieve the performance required for high-fidelity neural recordings [21–23]. Challenges such as suboptimal conductivity, electrode stability, and long-term biocompatibility have limited their widespread adoption, underscoring the need for further advancements in printed bioelectronics [24].

In this study, we present a low-temperature rapid prototyping approach that leverages photonic sintering to overcome critical limitations of printed neural interfaces. This process enables the fabrication of customizable, biodegradable and high-density electrocorticography (ECoG) arrays. By optimizing the combination of biocompatible gold nanoparticle inks and flexible polycaprolactone (PCL) substrates, we achieved a conformal, high-resolution neural interface. The developed ECoG device consists of an array of 36 gold electrodes, each 75 µm in diameter with a 350 µm pitch, spanning a compact 2.2 × 2.2 mm sensing area. To ensure reliable electrical performance, we implemented a dual-step post-processing technique involving oxygen plasma treatment followed by pulsed-light photonic sintering to enhance both conductivity and adhesion of the printed electrodes while maintaining substrate integrity. The fabricated arrays exhibited robust long-term stability, with average impedance values of 10.6 ± 4.5 kΩ at 1 kHz and minimal drift over a three-month period. Biocompatibility evaluations confirmed the absence of neuronal loss, inflammation, or gliosis throughout the implantation period. Functionally, the arrays successfully captured seizure propagation patterns in vivo in a rat seizure model with an SNR of 28 dB, among the highest reported for inkjet-printed ECoG devices of this scale. These findings position our platform as a scalable, high-performance solution for next-generation neural interfaces, offering a compelling route toward clinically viable, minimally invasive brain-machine interfaces.

## Results

### Low Temperature Additive Manufacturing of Biodegradable ECoG Arrays

Arrays of 36 gold 75µm-electrodes were successfully built on biodegradable PCL films via inkjet printing, covering an effective area of 2.2 x 2.2 mm^2^. A simplified schematic of the fabrication process is depicted in Figure 1a. Briefly, spin-coated PCL films were used as the backbone for the fully-inkjet-printed electrode array. Electrode fabrication was realized via inkjet printing using gold nanoparticles (AuNP) ink and epoxy-based UV-curable ink for insulation. The inkjet-printed electrodes were then sintered using a combination of photonic and plasmonic sintering. As shown in Figure 1b & c, the resulting device demonstrates remarkable flexibility and conformability, allowing it to seamlessly adhere to soft, curved surfaces, highlighting its suitability for interfacing with the brain.

**Figure 1.**
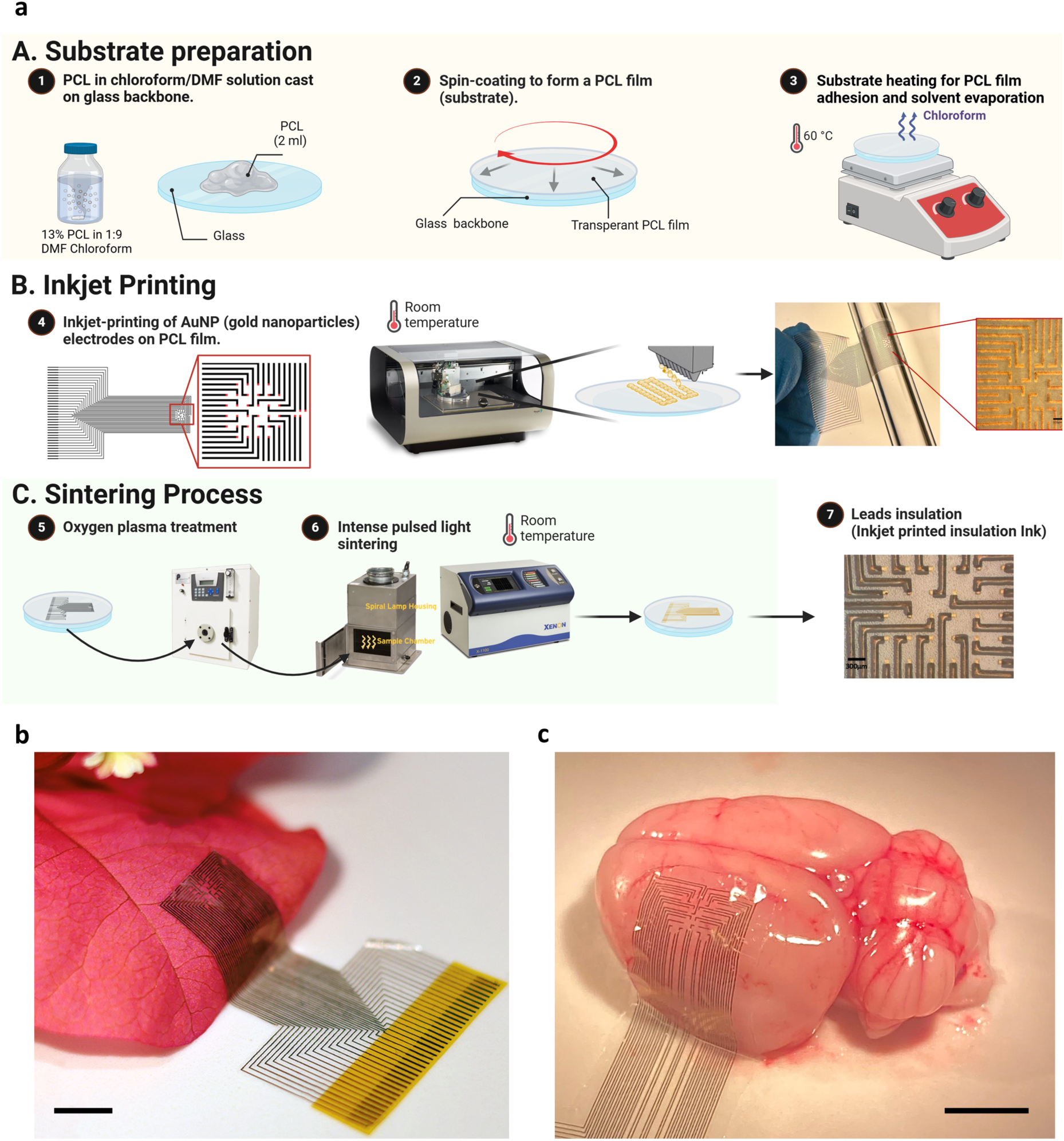
ECoG Array Fabrication. (a) Fabrication flow for inkjet-printed electrodes on PCL (polycaprolactone) film substrates. (A) Substrate preparation: (1) PCL in chloroform/DMF solution is cast onto a glass backbone, (2) spin-coated to form a transparent PCL film, and (3) heated to 60°C for adhesion and solvent evaporation. (B) Inkjet printing: AuNP (gold nanoparticle) electrodes are inkjet-printed onto the PCL film at room temperature to create patterned circuits. (C) Sintering process: (4) Oxygen plasma treatment prepares the surface, followed by (5) intense pulsed light sintering for conductivity at room temperature. (6) Insulation ink is inkjet-printed for lead protection, completing the fabrication process. This approach combines precision and scalability, enabling the creation of flexible, functional electronic components. (b) Inkjet-printed ECoG device placed on a delicate flower petal, demonstrating its ultra-thin and flexible properties suitable for conformal contact with soft, curved surfaces. Scale bar 5mm. (c) Device applied to an excised brain, illustrating its potential for neural interfacing applications with soft biological tissues. Scale bar 5mm.

Although layering in additive manufacturing can provide an advantage in improving performance (especially for conductivity), excessive layering can reduce printing resolution. Through iterative testing, we determined that three layers of gold nanoparticles provided uniform coverage with minimal spreading (Figure 2a). Optical and holographic imaging of the printed structures confirmed uniformity and integrity of the prints across the substrate (Figure 2b). Compared to other inkjet-printed ECoG arrays reported in the literature, our design achieves an electrode density among the highest reported, demonstrating significant advancement in spatial resolution and efficient substrate utilization [21, 25, 26].

**Figure 2.**
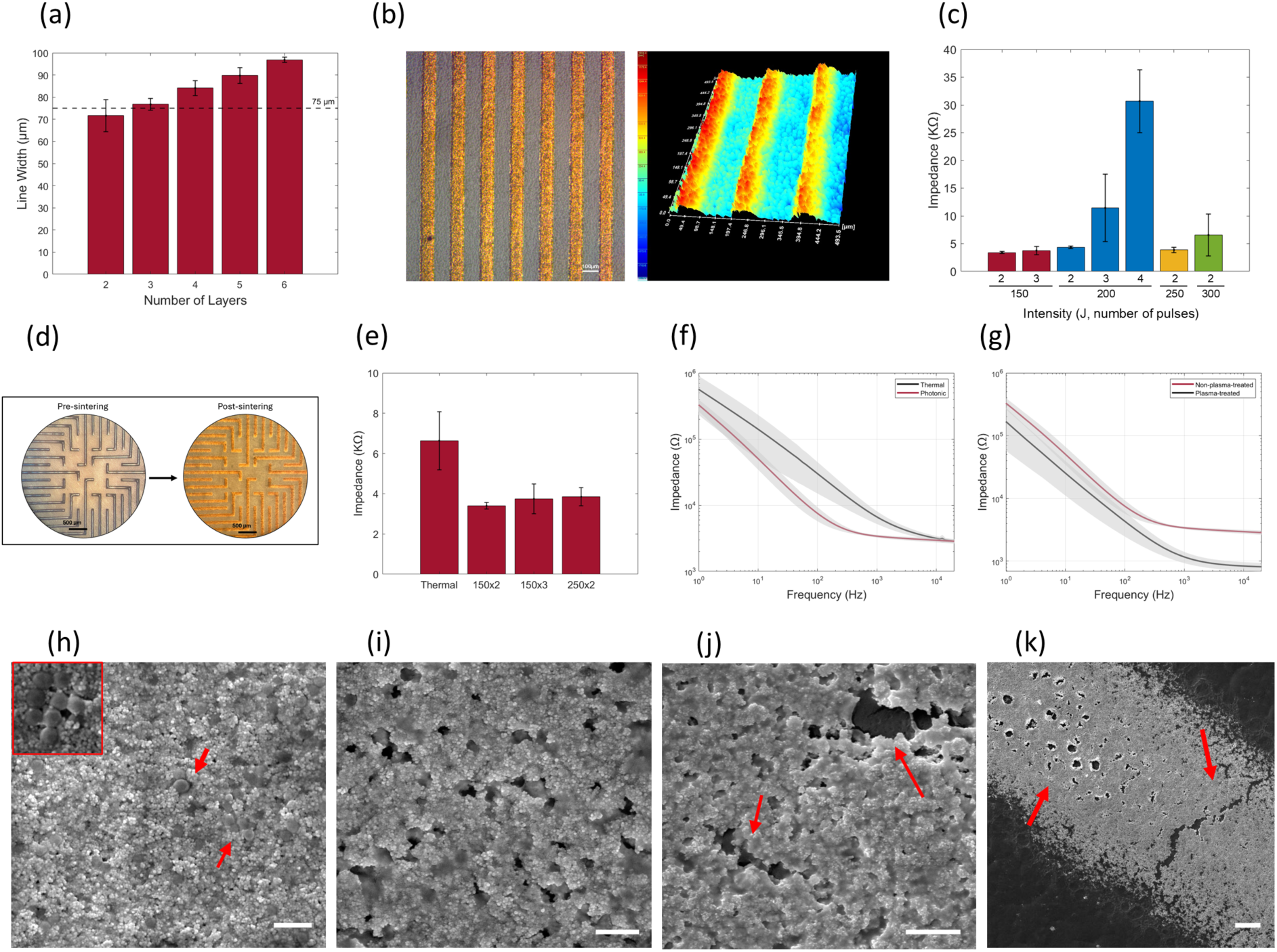
ECoG Array Characterization. (a) Variation of line width upon increasing number of printed gold nanoparticle layers (n=4). (b) Optical and DHM images of the 3D surface topography of printed gold-nanoparticle electrodes. (c) Electrochemical impedance at 1 KHz of 75µm electrodes sintered at energy levels ranging from 150 to 300 Joules at 2, 3 and 4 pulses. (d) Optical images of the electrode array before and after sintering, showing morphological and color changes post-sintering. (e) Impedance at 1 KHz of 75µm electrodes sintered via photonic sintering at diberent parameters and thermal sintering (n=3). (f) Ebect of thermal vs. photonic sintering (150J, 2 pulses) on the electrochemical impedance spectrogram of 75µm electrodes (n=3). (g) Ebect of 4-second plasma treatment pre-sintering on the electrochemical impedance spectrogram (n=3). SEM images of printed gold-nanoparticle structures (i) pre-sintering (j) after photonic sintering at optimal parameters (k) and (l) upon photonic sintering at higher intensity (350J, 2 pulses), revealing cracks and damage to PCL films (scale bars 500nm, 500nm, 1 µm and 10 µm respectively).

A major challenge in sintering gold nanoparticles on PCL substrates is its low melting point (55– 60°C), which restricts the use of conventional high-temperature sintering techniques. To address this, photonic sintering was investigated as a rapid, low-thermal-impact method to sinter gold nanoparticles while preserving the integrity of the PCL. Photonic sintering, via controlled intense pulsed light, induces rapid heating enabling the fusion of nanoparticles within milliseconds, limiting heat diffusion to the substrate. However, photonic sintering alone resulted in poor adhesion and occasional partial delamination of the gold structures from the PCL film. This led to inconsistency in electrode impedance across samples and the formation of microcracks at high energy levels. We therefore developed a two-step process combining oxygen plasma treatment with photonic sintering, which markedly enhanced the adhesion between the gold structures and the PCL substrate.

To optimize the curing process, photonic sintering parameters were adjusted based on electrode dimensions, pitch, and the gold electrode thickness. Dispersing the energy across multiple pulses, rather than a single pulse, was found to be advantageous to prevent sustained heating of the sample and minimized heat accumulation that leads to substrate damage. To determine the optimal parameters, pulse energy was systematically varied within the range of 150 to 300 Joules, delivered via 2-4 pulses for each energy level, with a 1-second inter-pulse interval. Energies below 150J failed to achieve curing, while energies exceeding 300J led to damage of the printed electrodes. Optimal conditions were identified at 150 J with two pulses, resulting in consistent electrode impedance of 3.4 ± 0.16 kΩ at 1 kHz (Figure 2c). We reported the impedance measurements at 1 kHz, as they are specifically designed for extracellular neuronal recordings, and this frequency closely corresponds to the spectral range of neuronal signal activity [27]. The visual transformation of the electrode array before and after photonic sintering, highlighted by a distinct color and texture change, is shown in Figure 2d, confirming morphological alterations associated with successful sintering. To further validate sintering efficacy, gold electrodes with similar dimensions were replicated on glass and were subjected to thermal sintering in an oven (180°C for 35 minutes). Electrochemical impedance measurements demonstrated that this alternative method of photonic sintering yielded superior outcomes at both low and high frequencies, highlighting its advantage in reducing thermal stress to PCL while enhancing overall electrode performance (Figure 2e & f).

To investigate the combination of plasma and photonic sintering, plasma durations ranging from 2 to 6 seconds were tested to identify the optimal treatment time that maximizes adhesion and reduced impedance variability without excessive etching of the gold layer. A 4-second plasma treatment was found to significantly reduce the average electrode impedance from 3.4 ± 0.16 kΩ to 1.16 ± 0.24 at 1 kHz (Figure 2g). Shorter treatment durations (<4 seconds) did not produce measurable improvements in impedance, whereas longer durations (>4 seconds) resulted in increased impedance (Supplementary figure 1a). This increase is likely attributable to surface damage caused by excessive etching of the gold electrodes’ surface.

The choice of different backbones, each with distinct heat dissipation properties, could play a crucial role in the sintering process and substrate integrity. In this work, we explored glass and aluminum as backbones for the PCL films. Electrodes on PCL films backed by aluminum required a total energy input of 1150 J to achieve comparable impedance values of 1–5 kΩ, whereas the more thermally efficient glass backbone required only 400 J delivered via a single pulse or 150 J via two pulses. A closer examination through SEM revealed visible damage to the PCL film in the aluminum-backed configuration (Supplementary figure 1b).

Nanoscale morphological changes in sintered gold electrodes were further examined via SEM imaging. Prior to sintering, gold nanoparticles displayed minimal aggregation and were surrounded by solvent particles (Figure 2h). After sintering, solvent particles were no longer visible, and the nanoparticles had coalesced, indicating improved conductivity (Figure 2i). However, sintering at higher-than-optimal energies induced micro- and macro-cracks, disrupting conductivity and damaging the PCL substrate due to heat conduction from the gold structures (Figure 2j & k). These findings emphasize the importance of precise energy control during photonic sintering to optimize electrode performance while maintaining substrate integrity.

### High-Resolution ECoG Recordings for Seizure Characterization

Prior to *in-vivo* experiments, the impedance profile of the electrodes was characterized, showing a steady decrease with frequency and an average impedance of 10.6 ± 4.5 kΩ at 1 kHz, with smaller deviations at higher frequencies (**Error! Reference source not found.**a). The phase angle peaked at mid-range frequencies, indicating capacitive behavior, and gradually shifted toward zero, reflecting dominant resistive behavior at higher frequencies (**Error! Reference source not found.**b). It is worth noting that electrodes with impedance below 600 kΩ at 1 kHz are deemed suitable for implantation [28–31]. Cyclic voltammetry (CV) confirmed the capacitive behavior of the electrode–electrolyte interface, as indicated by the characteristic rectangular current–voltage profile (**Error! Reference source not found.**c).

Figure 3d benchmarks our inkjet-printed ECoG array (square marker) against previously reported inkjet-printed flexible microelectrode arrays (circular markers). Our device achieved the highest electrode density (7.44 electrodes/mm2) of any device built with additive manufacturing with a compact electrode diameter of 75 µm, while maintaining a low average impedance of 10.6 kΩ at 1 kHz. By contrast, most existing devices trade off either density or impedance, with larger electrodes yielding lower impedance but lower spatial resolution. Our devices stand out by achieving both high spatial resolution and low impedance, a combination critical for neural recording.

**Figure 3.**
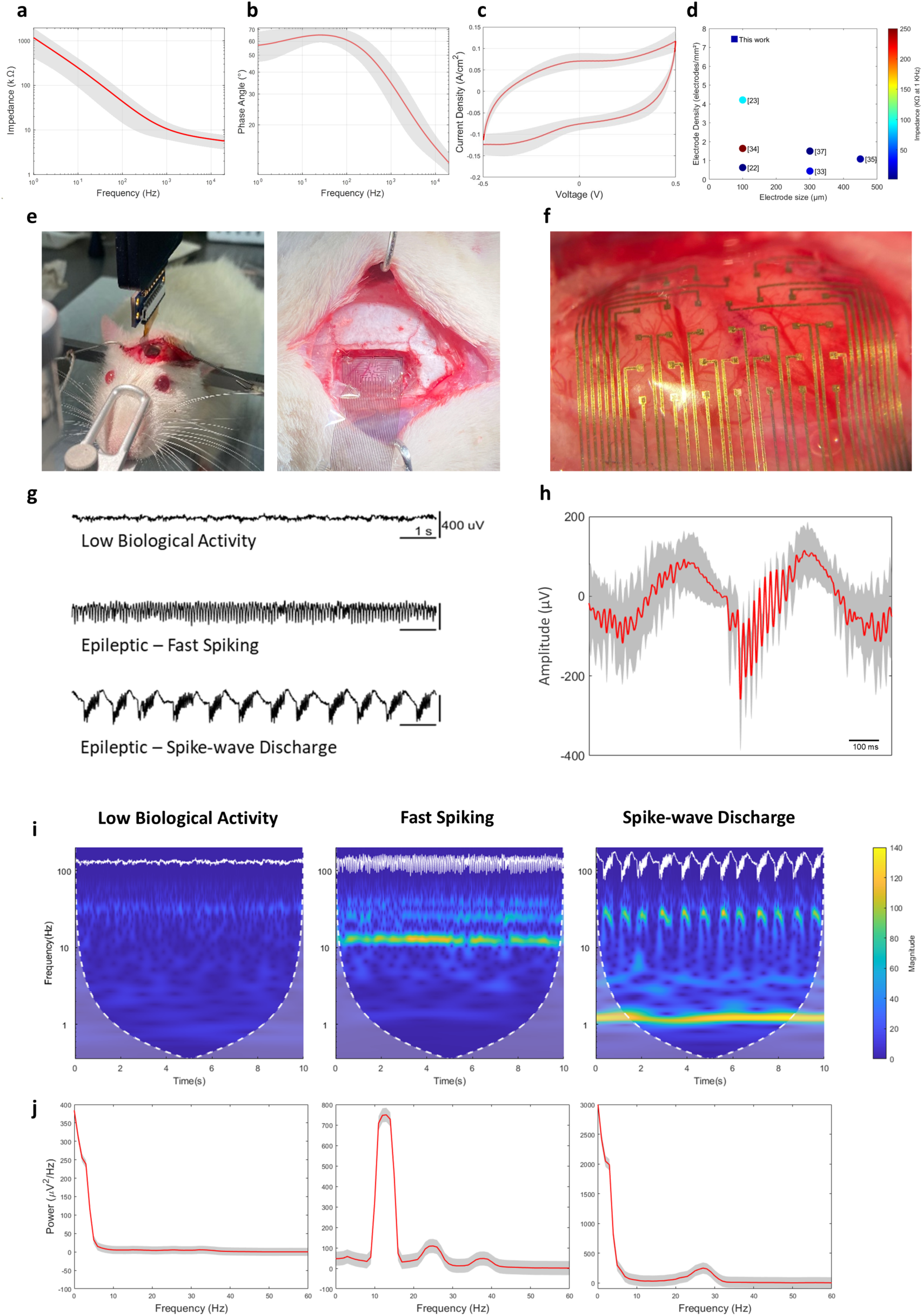
In vivo characterization. (a) Averaged electrochemical impedance, (b) phase angle and (c) cyclic voltammogram of 75 µm electrodes (n=36). (d) Comparison of the developed ECoG device against flexible electrode arrays fabricated via additive manufacturing in literature. Square data point represents our work (e) Device implantation for electrocorticographic recording from rat model. (f) Flexible ECoG array conforming to the cortical surface, demonstrating transparency and intimate contact with brain tissue. (g) Representative 10-second signals recorded during low biological activity (top), epileptic fast-spiking activity (middle) and epileptic spike-wave discharge (bottom). (h) Averaged recorded spike (red) from single electrode during spike-wave discharge 10 second interval. Standard deviation is represented as grey shaded region. (i) Scalograms of recorded ECoG signals during low biological activity (left), epileptic fast spiking (middle), and epileptic spike-wave discharge (right) and (j) corresponding averaged power

To asses *in vivo* performance, the arrays were implanted in adult male Sprague-Dawley rats (n=4), weighing between 300-350 g. Following the placement of the electrode array onto the cortical surface, successful electrocorticographic recordings were obtained under anesthesia and during epileptic seizure (**Error! Reference source not found.**e). The high flexibility of the OECT array allowed it to conform seamlessly to the brain’s surface, minimizing the risk of trauma and ensuring stable contact with the underlying tissue, thus contributing to high signal quality (**Error! Reference source not found.**f). The device’s transparency is also evident from the clear visualization of vasculature on the cerebral cortex. After seizure induction via intra-amygdalar kainic acid (KA) injection, electrocorticographic recordings were collected simultaneously from 32 electrodes over one hour. The KA injection induced status epilepticus with an electrographic onset of around 20 minutes post-injection. The status epilepticus was characterized by recurrent seizures without a return to baseline activity throughout the one-hour recording period. Recordings revealed three spatiotemporally distinct patterns of neural activity that were consistently observed across all experiments (**Error! Reference source not found.**g): (1) minimal, irregular biological activity under anesthesia, characterized by a low-amplitude, slow signal, reflecting a baseline or quiescent brain state; (2) epileptic fast spiking activity indicating the onset of epileptiform activity, and the brain’s transition towards a seizure state; (3) epileptic spike-wave discharge, characterized by synchronized, pronounced spike and wave patterns, a hallmark of an evolving ictal phase, as recorded in the monitored hemisphere [31–33]. A representative averaged spike waveform recorded by a single channel during the spike-wave discharge phase illustrates the consistent shape and timing of the spike and wave components across multiple events, with minimal variation as indicated by the narrow standard deviation envelope (**Error! Reference source not found.**h). Following the spike-wave discharge phase, postictal suppression was observed, marked by an amplitude lower than the baseline state, suggesting transient cortical silencing immediately after seizure termination. Notably, both fast-spiking and spike-wave discharge appeared in alternating succession with recurrent bursts and were consistently observed in all recorded seizures within the recorded cortical region, indicative of focal seizure dynamics with local network propagation. Our ECoG array enabled sensitive detection of subtle electrophysiological changes across distinct stages of epileptic seizures. The average SNR of the obtained recordings during the spike-wave discharge phase was found to be 28 dB, the highest reported for inkjet-printed ECoG electrodes of this size and density (Table 1).

**Table 1.**
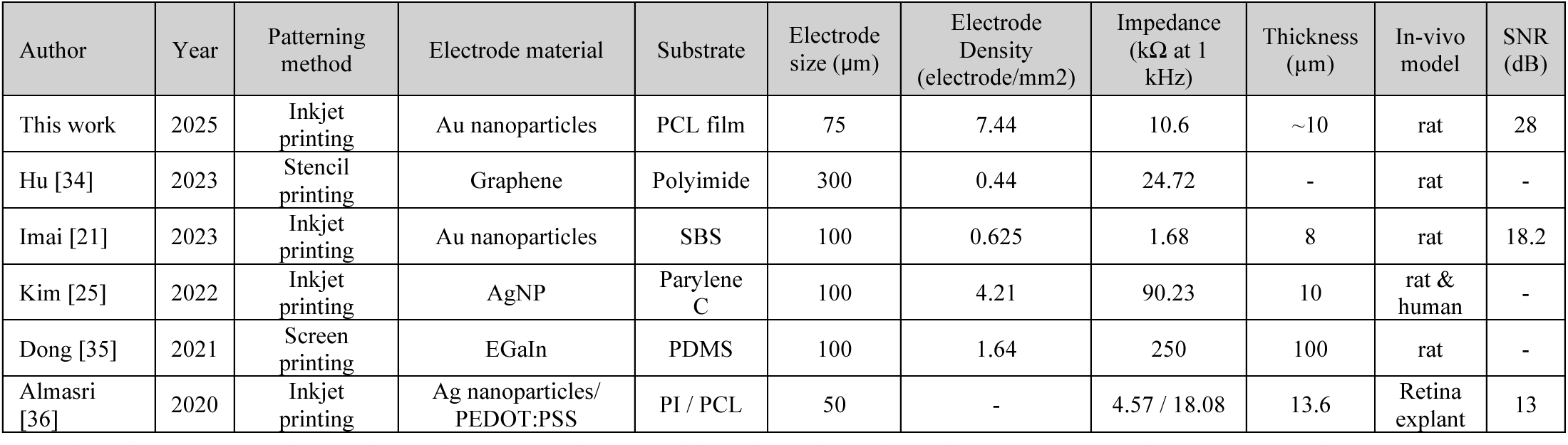
Properties of flexible microelectrode arrays for electrophysiological monitoring developed using additive manufacturing techniques (for devices with diameter ≤ 300 µm).

To further analyze the electrocorticographic signals, time-frequency scalograms and average power spectra for 10-second sections of normal and epileptic brain activity were generated (**Error! Reference source not found.**i & j). The scalograms clearly distinguish between the distinct types of activity, providing a detailed representation of the temporal and frequency dynamics. It was noted that the power spectra during seizure status exhibited greater amplitude across the entire frequency range compared to the normal state, especially in the low-frequency band (<40 Hz), indicating heightened overall neuronal activity. During periods of low biological activity, the scalogram showed minimal power across all frequencies. As the brain activity progresses into the fast-spiking phase, the scalogram revealed a notable increase in power at higher frequencies, particularly around 13 Hz, corresponding to the fast spiking seen in the time-domain recordings, with lower peaks observed at around 25 and 38 Hz. This increase in power confirms the presence of rapid, synchronous neuronal firing indicative of the onset of epileptiform activity. During the spike-wave discharge phase, two distinct spectral bands were observed: one below 2–4 Hz, likely reflecting the repolarization phase following sharp spikes (i.e., the slow spike wave component), and another around 27 Hz, typically associated with fast spiking activity. This separation suggests the presence of two neuronal populations: one contributing to the slow spike–wave pattern, and another generating potentially low-amplitude, high-frequency spiking superimposed on this slower activity. While such dual-frequency components are not commonly reported, the high spatiotemporal resolution of our recordings may be capturing distinct spiking dynamics not previously observed.

### Machine Learning Model for Neural Activity Classification

To demonstrate the translational potential of our neural interface, we implemented a deep learning–based seizure classification model on ECoG signals recorded using the inkjet-printed arrays. A one dimensional (1D) convolutional neural network (CNN) was trained using ECoG recordings from two selected channels (E14 and E31) obtained during status epilepticus. The signals were segmented into 2-second windows and manually annotated into three distinct neural activity classes: baseline (green), fast spiking (blue), and spike-wave discharges (red). Representative signal epochs for each class are illustrated in Figure 4a.

**Figure 4.**
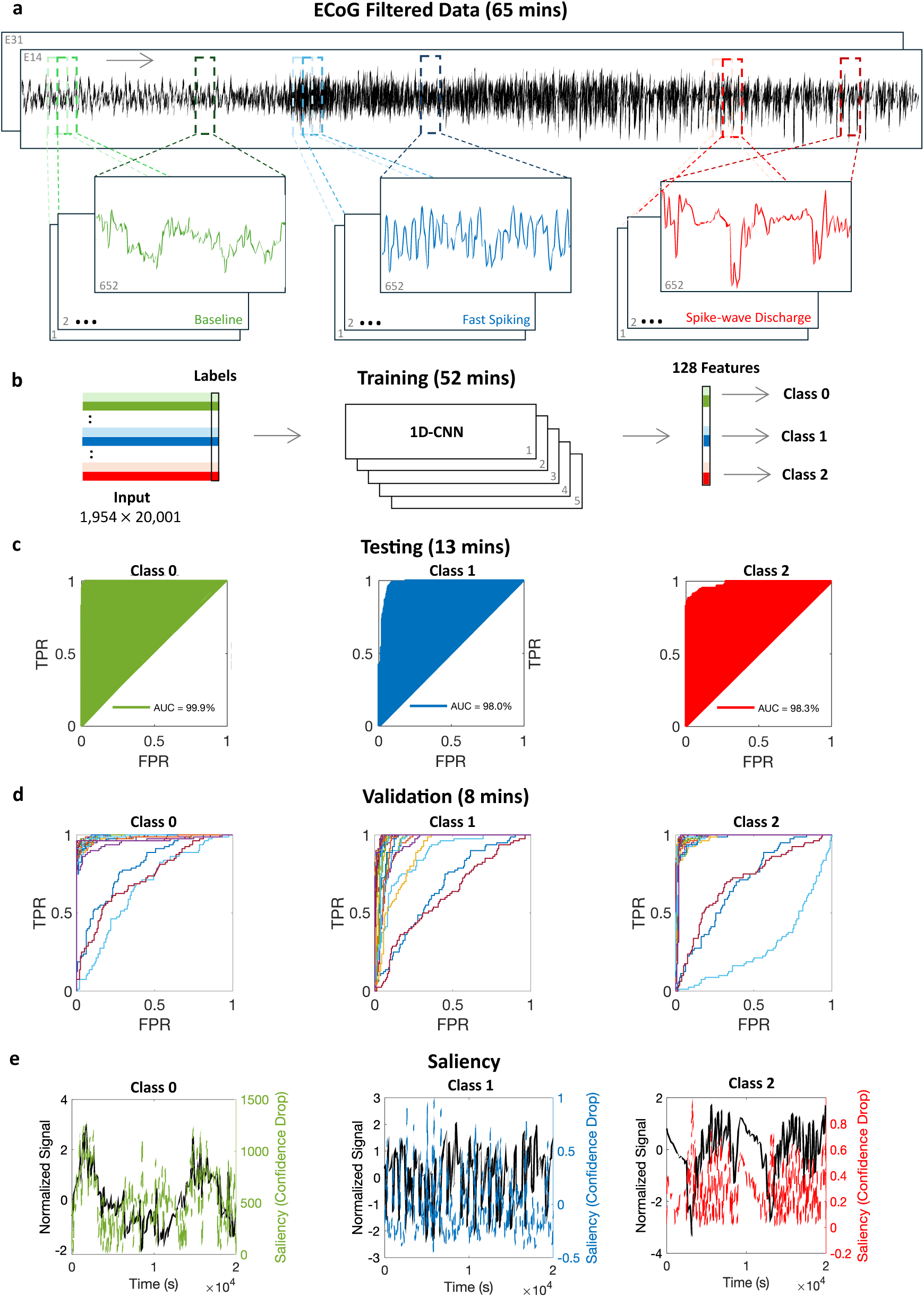
Classification of epileptic activity using a 1D-CNN. Classification of distinct neural activity stages reflecting seizure evolution using deep learning. (a) Filtered ECoG signals from two trials (channels E14 and E31) spanning 65 mins were segmented using a 2-second sliding window with 50% overlap. Labels were assigned to each segment based on expert annotations into three classes: Class 0 (baseline), Class 1 (fast spiking), and Class 2 (spike-wave discharge). (b) A total of 1,954 × 20,001 samples, equally distributed across the classes, were used to train 27-layer 1D-CNN model. (c) Testing on 13 mins of ECoG data demonstrated high classification performance: 99.9% accuracy for Class 0, 98.0% for Class 1, and 98.3% for Class 2. (d) Validation over 8 mins of signals from all channels confirmed generalizability, with consistently high metrics except for three lower-performing channels (E13, E15, and E21). (e) Saliency maps highlight the temporal regions within the ECoG signal that most contributed to classification decisions, confirming biologically meaningful patterns for each seizure type.

In total, 1,954 annotated 2-second segments, each containing 20,000 time points, were used for training and testing. After hyperparameter tuning, the best performing (kernel sizes [9,7,5], L2 regularization λ = 0.001) achieved a validation accuracy of 92.95%. Successive convolutional layers (32, 64, 128, 256, 256 filters) extracted progressively higher-order temporal features from the raw ECoG input, which were then compressed into a 128-dimensional latent feature space before final classification (Figure 4b).

On the held-out test set, the model achieved an overall accuracy of 93.1% with a precision of 93.8% and recall of 93.1%. Class 0 (baseline) was separated with near-perfect AUC of 99.9%, correctly identifying 123/130 non-seizure segments (Figure 4c, green). Importantly, no seizure events were misclassified as baseline, demonstrating clear separation between pathological and non-pathological activity. For seizure subtypes, performance remained high (AUC = 98.0% for Class 1 and 98.3% for Class 2), confirming strong predictive performance across seizure subtypes. Performance was slightly lower for these classes (Figure 4c), with minor misclassification occurring between fast spiking (blue) and spike-wave discharge (red), which is expected biologically as fast spiking often precedes and transitions into spike-wave discharges in rodent models, leading to temporal overlap at seizure onset [37, 38].

Beyond three-class task, we assessed binary and combined discrimination. When Class 0 (baseline) was compared with the combined seizure classes (1 and 2), the model achieved 97.7% accuracy with an AUC of 99.4%, confirming its potential for real-time detection. Among binary tasks, Class 0 vs. 2 yielded the highest performance (99.6% accuracy, AUC = 100%), with complete separation of baseline from spike-wave discharge due to their distinct morphology (only a single spike-wave segment was misclassified). In contrast, Class 0 vs. 1 was more challenging (95.8% accuracy, AUC = 99.9%), as fast spiking may exhibit spectral similarities with baseline high-frequency activity.

Collectively, these results demonstrate that while seizure detection is near-perfect, subtype discrimination is influenced by the natural progression of seizure dynamics and the inherent biological overlap between transitional neural states, rather than reflecting a limitation of the model. Detailed confusion matrices and additional metrics for the three-class, binary, and combined classification tasks are provided in the supplementary material. Future work may integrate recurrent neural networks (RNN) to better capture evolving seizure dynamics over longer timescales.

To assess generalizability to new recordings, the three-class discrimination model was further validated on an 8-minute independent ECoG recording obtained with the flexible chip and unseen during training. The dataset included all 32 channels, segmented into 2-second windows. The model achieved an average accuracy of 85.6%, recall of 85.6%, precision of 87.0%, F1 score of 84.3%, and average AUC of 95.4%. These results indicate robust and balanced performance in a continuous, real-world-like setting. Specifically, channels E1 and E12 showed near-perfect AUCs (>99%), while E5, E6, E9, E11, E14, and E25 maintained AUCs above 98%. In contrast, E13 (AUC = 60.6%), E15 (AUC = 72.1%) and E21 (AUC = 68.5%) exhibited the lowest classification performance, likely due to reduced SNR. A complete table of per-channel performance is provided in the supplementary material. The classifier’s multi-class performance is shown in Figure 4d, illustrating consistent classification performance and high reliability across varied data channels.

For interpretability, saliency maps were computed from the training set to visualize class-relevant contributions along the input signals. The original z-score normalized signal (black) was overlaid with class-specific relevance confidence drop scores, highlighting which regions most influenced model predictions as shown in Figure 4e. For Class 0, saliency aligned with low-amplitude and stable baseline patterns (Figure 4e, green). For Class 1, attention was directed toward short, irregular, and rapid fluctuations typical of fast spiking (Figure 4e, blue). In Class 2, saliency focused on rhythmic activity patterns and high amplitude segments of spike-wave discharge (Figure 4e, red). These maps confirm that the model learns to associate time-domain morphology with biologically distinct phases of seizure evolution.

### Spatiotemporal Mapping of Seizure Propagation

Simultaneous 32-channel recordings were used to generate topographic pseudocolormaps, providing a visual representation of cortical activity (Figure 5a). Sequential colormaps generated at a frame interval of 50ms during a 6.4-second section of spike-wave discharge demonstrated our device’s efficacy in accurately localizing seizure onset regions and tracking seizure progression with high spatiotemporal resolution (Figure 5b, Supplementary Video). To assess reproducibility, topographic maps were generated from six consecutive seizures during the spike-wave discharge phase (Figure 5c). each at the peak with the highest amplitude. These colormaps revealed a consistent spiral propagation pattern across consecutive seizures, demonstrating the array’s capability for high-resolution cortical mapping and dynamic seizure monitoring. This suggests a consistent pattern of cortical recruitment that may reflect the involvement of specific adjacent neuronal networks in repeated activation sequences.

**Figure 5.**
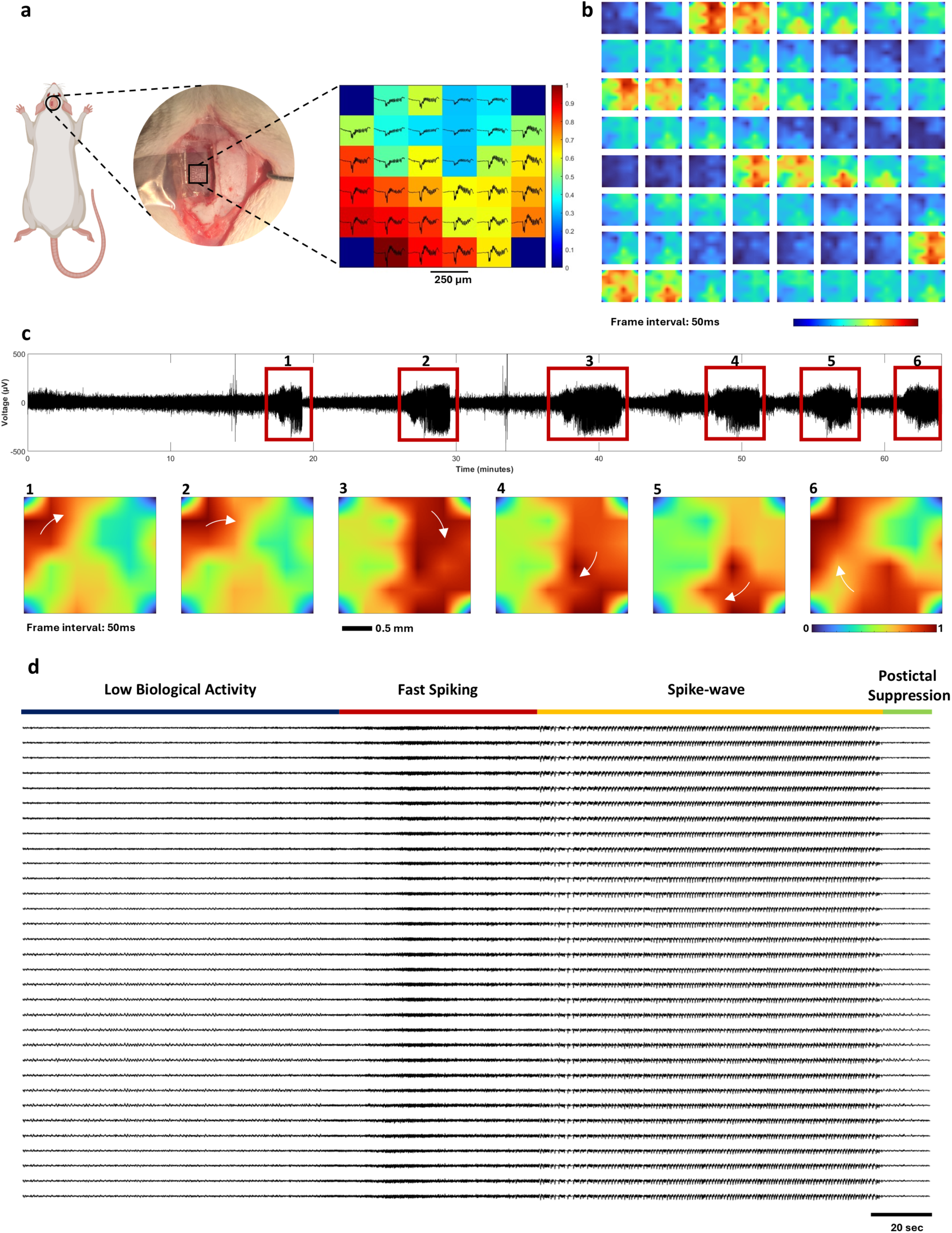
Cortical mapping for seizure monitoring. (a) Placement of the ECoG device on the cortical surface of a rat, with an inset showing the heat map generated. The heat map illustrates the spatial distribution of neural activity. (b) Sequential topographic colormaps of 32 electrodes at frame interval of 50 ms during a 6.4-second section of epileptic spike-wave discharge (c) 1-hour ECoG signal from single electrode and corresponding sequential topographic color maps from 32 recording sites of six consecutive seizures. (d) Representative 32-channel ECoG traces (5 minutes) showing the temporal progression of seizure activity for a single seizure.

Examination of the simultaneous 32-channel recordings reveals a predominantly synchronous progression of seizure activity across the covered cortical surface (Figure 5d). The transition through distinct phases is evident across all channels, indicating widespread network involvement during the ictal event. However, subtle variations in onset timing and signal amplitude were observed, particularly during the transition into the fast-spiking and the spike-wave discharge phase. These variations likely reflect spatial propagation of epileptiform activity across the cortex, with certain regions initiating or amplifying activity earlier than others. The detection of these subtle variations, enabled by the high spatial and temporal resolution of our array, could offer valuable insight into seizure onset dynamics and cortical activation patterns during ictal events.

### Mechanical Testing

Upon subjecting the developed arrays to mechanical stresses, electrode impedance increased by 12% after 100 cycles of bending and by 9% after 100 cycles of twisting (Figure 6a & b). It is important to note that these cycles represent an extreme case of mechanical stress not typically encountered in acute experiments. The observed increase in impedance, though measurable, is relatively slight and suggests that the electrodes maintain their functional integrity despite the rigorous testing. The slight impedance rise likely reflects microcrack formation in conductive leads, a common effect of repeated stress in flexible electronics.

**Figure 6.**
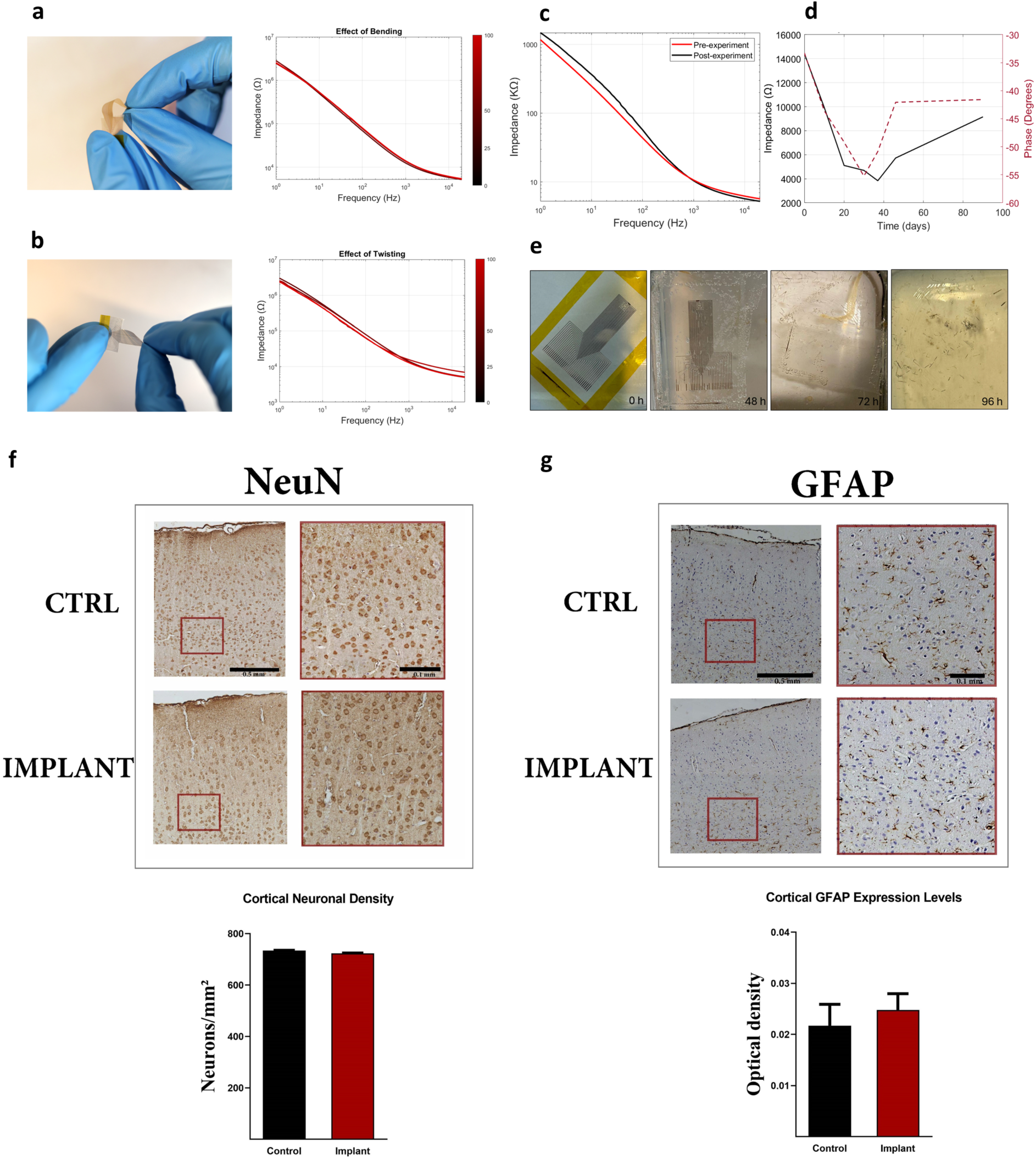
Device stability and biocompatibility. Ebect of (a) bending and (b) twisting (100 cycles) on electrical performance of ECoG electrodes. (c) Average impedance of 36 electrodes before and after electrocorticography experiment. (d) Stability of impedance and phase angle measurements at 1 KHz over 3 months. (e) Gradual accelerated degradation of the PCL-based device with gold electrodes over time. Comparison of (f) cortical neuronal densities and (g) astrocytosis between the control and implant groups. NeuN staining reveals no significant diberence in neuronal density between the control and implant

### Device Stability

To evaluate the stability of our device, electrochemical impedance of the electrodes (n=36) was measured before and after experiments to evaluate any potential degradation or changes in functionality due to the experimental stresses (Figure 6c). Notably, the impedance at 1 kHz remained consistent throughout. Such stability is crucial for ensuring reliable and reproducible data acquisition in neural applications. Additionally, the long-term stability of our device was also assessed by the continuous measurement of the electrodes’ impedance at 1 kHz every hour over a three-months period (Figure 6d). Initially, both the impedance and phase angle measurements showed a downward trend, stabilizing after around 30 days. Subsequently, a slight increase was observed, suggesting a minor reduction in the electrode performance. Despite these variations, the overall trend shows that the impedance remains within a consistent range of 3 to 14 kΩ at 1kHz indicating robust stability of the electrode array over extended periods. Similarly, the phase angle demonstrated relative stability, fluctuating between 33° and 57°. Beyond its recording capabilities, the device was designed to be fully bioresorbable, eliminating the need for secondary removal surgeries. An accelerated degradation test was conducted to assess the biodegradation of the PCL-based ECoG device. The device was fully immersed in 10 ml of phosphate-buffered saline (PBS) with a pH of 13 at 37°C to simulate conditions that promote degradation. The degradation process was monitored over a period of 4 days, with the results shown in Figure 6e. At the 48-hour mark, the gold electrodes printed on the PCL substrate began to detach, marking the initial stage of degradation. By 72 hours, the gold electrode prints had completely disappeared, while the PCL film remained largely intact. However, after 96 hours, the PCL substrate fully disintegrated into small fragments. This degradation process was driven solely by chemical hydrolysis, as no enzymatic activity was involved under these test conditions. The presence of enzymes in vivo would likely accelerate the process, further facilitating the breakdown of the device. In the body, various factors contribute to the degradation of implantable devices, including hydrolysis, oxidation due to reactive oxygen species produced by tissues, physical degradation due to swelling and mechanical loading as well as wearing, and enzymatic degradation [36, 39].

### Biocompatibility

To evaluate the biocompatibility of our ECoG device, we assessed neuronal integrity and glial response in the cortex following implantation. Specifically, we examined neuronal loss and astrocytosis by performing immunohistochemical staining for NeuN and GFAP on cortical tissue sections. The results, as shown in Figure 6f & g, revealed that both the neuronal density and the GFAP levels were comparable between the implant group and the control group, suggesting that the ECoG device does not induce chronic injury in the form of overt neuronal damage or reactive astrogliosis in the cortex. This is a promising indication of the device’s potential for long-term implantation and use in neural recording applications.

## Discussion

The neural interface developed in this work represents a significant step forward in designing high density, low-impedance electrode arrays. This fully inkjet-printed neural interface demonstrates a unique combination of small electrode size, high electrode density, and compact sensing area with robust electrical performance on a flexible substrate [36]. The use of biodegradable polycaprolactone (PCL), considerably softer than conventional substrates such as silicon, polyimide, or parylene-C, improved mechanical conformity to the brain surface while preserving the device’s structural integrity. The fabrication process is highly efficient, enabling device production in under 24 hours at a substantially lower cost than conventional methods, making it accessible for a broader range of applications. Mechanical testing and long-term impedance monitoring demonstrated the device’s robustness -an essential attribute for neural interfaces intended for prolonged use in both clinical and research settings. The PCL-based arrays completely degraded under simulated physiological conditions, and biocompatibility evaluations revealed no significant neuronal loss or glial response, affirming the device’s safety for neural interfacing.

By integrating plasma pre-treatment with photonic sintering, this study developed a sintering strategy that simultaneously optimized conductivity, adhesion, and overall functionality. This synergistic approach addresses the fundamental challenges of sintering on thermally sensitive substrates, marking an important advance in the fabrication of flexible bioelectronics.

Conventional ECoG electrodes typically have a diameter of approximately 4 mm and an interelectrode distance of about 10 mm [40]. This relatively large size limits spatial resolution and hinders precise localization of seizure foci, contributing to poor surgical outcomes in up to 50% of patients [41]. Our inkjet-printed ECoG array offers a significant improvement in spatial resolution and recording fidelity. The device successfully captured distinct phases of seizure activity in vivo with a signal-to-noise ratio of 28 dB, outperforming previously reported devices of similar scale and fabrication techniques.

The 1D-CNN model we developed classified time-series signals recorded directly from the implanted ECoG array without frequency-domain transforms or extensive preprocessing. While frequency-domain analyses have historically dominated this field, time-domain features have recently shown sufficient discriminatory power to differentiate complex neural states [42]. Barry et al employed CNN-based cross-patient seizure classifiers on large-scale human ECoG datasets by converting time-series ECoG data into spectrograms, achieving up to 95.7% accuracy [43] . Similarly, the artificial neural network system developed for single- and multi-channel intracranial EEG in rodent models reported >98% sensitivity across 2892 h of recordings using manually engineered features [44]. In comparison, our combined seizure vs. baseline model trained directly on raw ECoG time-series achieved 97.7% accuracy with a sensitivity of 97.9%, despite being trained on a smaller dataset with shorter recordings. Training with only two channels was sufficient for effective classification, as these channels were among the most informative in terms of signal quality (low vs. high SNR). This not only reduces computational complexity but also enhances the feasibility of integration into real-time or embedded systems. Previous studies have also demonstrated the efficacy of low-channel seizure detection models, where two-channel setups achieved performance comparable to high-density systems [45]. This supports our design choice to employ 1D-CNN on raw ECoG signals, validating the ability of temporal CNNs to capture the seizure dynamics while preserving the physiological integrity of the signals. The fabricated device provided high-resolution recordings and real-time access to neural activity, with flexible switching between recording and inference. Together, these features make CNN-based classification well-suited for real-time, embedded seizure monitoring. Beyond binary classification, the model captured complex temporal dynamics across seizure stages, providing a basis for phase-specific detection, thus improving diagnostics. This capability is particularly valuable in online monitoring systems, where identifying pre-ictal fast spiking or transitional activity could enable timely intervention before full spike-wave discharges onset.

Overall, our deep learning-based model achieved consistent performance across both test and validation datasets, comparable to state-of-the-art approaches. More importantly, it demonstrates the translational potential of our flexible ECoG array in real-time neural monitoring. This work lays the groundwork for closed-loop epilepsy management systems, where flexible electrode arrays and lightweight neural models enable timely, interpretable, and reliable seizure detection. Future enhancements may include adaptive windowing, hybrid temporal-spectral features and expanded datasets.

Colormaps generated from simultaneous 32-channel recordings demonstrated the capability of our device to precisely identify the seizure zones and monitor the propagation of seizure activity in real-time. Notably, while spiral wave propagation during seizures has been observed in prior studies, particularly in animal models and computational simulations, our findings reveal a consistent spiral-like pattern across multiple seizures within the recorded region [46–49]. This pattern suggests that certain cortical regions may serve as hubs for seizure propagation, potentially due to intrinsic network properties or localized energy dynamics. The high spatiotemporal resolution of our array enabled the detection of these subtle yet consistent patterns, offering insights into seizure perpetuation mechanisms during status epilepticus. Importantly, this spiral motif was identified within a small, specific region of the cortex, raising the possibility that similar patterns may emerge elsewhere across other cortical areas during seizure spread. However, given the limited spatial coverage of our current array, we cannot confirm whether these motifs extend across the cortex or represent isolated local phenomena. This underscores the need for future studies using larger-scale, high-density ECoG arrays capable of mapping cortical activity over broader areas while preserving high spatiotemporal resolution.

The precision achieved by our inkjet-printed ECoG array is crucial not only for improving seizure detection and classification but also for advancing real-time monitoring in clinical settings. Thus, beyond epilepsy applications, the high-resolution mapping capability of our device has broader implications for neurotechnology, particularly in brain-computer interfaces. This technology could be leveraged to detect and decode high-frequency neural activity patterns associated with motor and cognitive processes, making it a potential tool for decoding movement intent in BCIs with micrometer-scale resolution, surpassing the capabilities of existing devices [50]. This aligns with previous work demonstrating that ECoG-based BCIs can classify imagined movements with up to 80% accuracy using motor cortex signals [51]. Our device offers the resolution and stability needed to further advance these outcomes.

## Conclusion

This work demonstrates a new class of electrocorticography (ECoG) arrays that combines biodegradability, rapid prototyping, and high-performance neural recording within a single scalable platform. By leveraging additive inkjet printing and photonic sintering, we achieve rapid fabrication of high-density gold nanoparticle electrodes on flexible polycaprolactone substrates, eliminating the need for traditional cleanroom microfabrication. In vivo recordings enabled precise seizure mapping, while histology confirmed minimal inflammation after chronic implantation. Integration with a deep-learning model yielded over 95% accuracy in seizure stage classification, demonstrating the potential for real-time, closed-loop neuromodulation. This additive-manufacturing approach offers a scalable, customizable, and environmentally sustainable route for next-generation neural interfaces that are disposable, patient-specific, and clinically translatable.

## Methods

### ECoG array design and fabrication

**Substrate.** PCL solution of 13% w/v PCL in 9:1 chloroform to dimethylformamide (DMF) was prepared by first dissolving PCL pellets (704105 Sigma-Aldrich) in chloroform by magnetic stirring at 50°C for 30 minutes. Following complete dissolution, dimethylformamide (DMF) was added to the solution and stirred for 10 minutes at room temperature. PCL films were then prepared by spin coating PCL on clean glass slides at 600 rpm for 40 seconds to form thin films of a uniform thickness of 10 μm. The slides were then placed on a hotplate at 60°C for 1 minute, enhancing adhesion between the PCL film and the glass.

**Inkjet printing.** For electrode fabrication, a Dimatix DMP-2850 inkjet printer (Fujifilm Dimatix, USA) fitted with piezo-driven 12-nozzle printheads with a drop volume of 2.4 pL was employed [20]. A 36-electrode array design with leads and contact pads was printed on the PCL substrate by depositing gold nanoparticles (AuNP) ink (JG-125 Novacentrix, USA). Each electrode in the array was designed with a surface area of 75×75 µm². The leads were fabricated with a consistent width of 75 µm to maintain signal integrity while accommodating the entire array within a compact footprint of approximately 2.2 × 2.2 mm^2^. As specified by the adapter (Multi channel systems connector ADPT-FM-32) used for device testing, the first and second leads on each side were designated as the ground and reference connections, respectively.

No further modification was required for the ink since it was compatible with the printer utilized. The printing drop spacing was chosen to be 25 µm for optimal printing quality. The number of printed layers was varied from 2 to 6 layers to determine the optimal layer count that minimized spreading while maximizing electrical performance.

**Insulation.** Inkjet printing was used to insulate the conductive leads using a epoxy-based UV-curable ink (DM-INI-7003-Dycotec Materials, UK), ensuring the conductive pathways were protected while leaving the 75 µm electrode sites and connector pads exposed for electrical interfacing. This UV-curable ink was applied with high precision, covering leads with a width of 125 µm, while the rest of the array was insulated using a PCL film to maintain necessary exposure. The UV-curable ink constituted 0.63% of the device material and had been previously tested for biocompatibility. Dielectric testing was conducted to determine the number of printed layers required for optimal insulation, establishing that four layers of the UV-curable ink were necessary.

**Photonic sintering.** Inkjet-printed samples were sintered using the room temperature Xenon X-1100 High-Intensity Pulsed Light (IPL) system, chosen for its compatibility with the heat-sensitive PCL substrate. The photonic system delivers up to 9 J/cm² of radiant energy per pulse, with pulse durations adjustable from microseconds to milliseconds. For this work, pulse durations of a few milliseconds, energy levels of 1–2.5 J/cm², and pulse frequencies of 1–3 Hz were employed. The number of pulses was optimized based on the thickness, substrate properties, and pattern configuration of the array. Samples were positioned at a uniform distance of 6.5 cm beneath the LH-912 lamp housing to ensure consistent exposure. A test pulse was performed to optimize settings, with adjustments made as necessary [52, 53]. Prior to sintering, oxygen plasma treatment (E-25 model, PlasmaEtch) was employed to ensure complete rapid drying of the samples, thus reducing total device fabrication time. Plasma treatment was done at 499.3 mTorr, 150 W, 15 cc/min oxygen flow rate for 2 to 6 seconds [54].[54]. Post-sintering inspections verified quality and electrical conductivity.

### Device characterization

**Electrochemical Impedance and Cyclic Voltammetry Analysis.** A four-point probe system (Ossila, UK) was used to study the effect of the number of layers and sintering parameters on the sheet resistance of the printed gold structures. Electrochemical characterization of the electrodes was performed via a Potentiostat/ Galvanostat/ZRA system (Gamry Instruments, USA) using a three-electrode cell, where the electrochemical impedance spectroscopy (EIS) and cyclic voltammetry (CV) were employed to measure the impedance and charge injection capacity of the electrodes. The charge storage capacity (CSC) was subsequently inferred via calculating the area under the CV curve (swept between −0.5 and 0.5 V at 1V/s). Phosphate buffer saline (PBS) was used as the electrolyte medium; a platinum (Pt) electrode was used as a counter electrode while Ag/AgCl wire was used as a reference electrode.

**Scanning Electron Microscopy** Scanning electron microscopy (SEM; MIRA 3 LMU Tescan, Czech Republic) was employed to assess the surface morphology of the fabricated electrodes. The primary focus was to observe how different sintering parameters affected the structural integrity of the nanoparticles. SEM imaging provided high-resolution insights into the changes in the array after sintering, enabling a detailed evaluation of the nanoparticle fusion and the overall quality of the sintered array, crucial for optimizing electrode performance [55, 56].[55, 56].

**Mechanical Testing.** To evaluate the impact of mechanical stresses on the performance of the ECoG array, the developed devices were subjected to 100 cycles of bending (90°) and twisting (360°), and their electrochemical impedance spectrograms were obtained after every 25 cycles.

### In-vivo assessment

**Animal experiment.** All experiments were approved by the Institutional Animal Care and Use Committee (IACUC) at the American University of Beirut. The rats were first anesthetized via intraperitoneal injection of Ketamine and Xylazine (80/20 mg/Kg). The fur from the area between the eyes to the ears was shaved, and the animals were mounted on a stereotaxic apparatus using ear bars placed in the auditory meatuses and a mouthpiece set at the appropriate height for adult rats. The head was scrubbed with isopropyl alcohol and betadine. A midline incision was made using a scalpel, and clamps were used to retract the skin, exposing the skull. Underlying tissues were cleaned using a sterile swab, revealing the bregma and lambda. Coordinates of the bregma were identified, and the edges of a 4×4 mm craniotomy site were accordingly marked on the left side of the brain above the somatosensory area. A surgical drill was used to remove the bone, followed by the removal of the dura using fine-tip forceps. For seizure induction, a 1µL Hamilton syringe fixed to a stereotaxic holder was used to inject 0.6µL of 1mg/ml kainic acid (KA) in saline into the amygdala (−2.8mm AP, 5mm ML, and 8.8mm DV from the bregma). Following the injection, the electrode array was placed on the surface of the cortex and connected to the ME2100 data acquisition system (Multichannel Systems, Germany), to record data from 32 channels simultaneously at a sampling frequency of 10 kHz with the integrated analog band-pass filter set from 1-to-500Hz over a period of 1-hour post-KA injection.

**Data Analysis.** The ECoG data were processed offline using a 200Hz digital lowpass filter and a 50Hz notch filter. Scalograms were generated from 0 to 200Hz over the period of 10 seconds using the Morlet wavelet. The signal to noise ratio (SNR) was calculated as the difference between the decibel power of the highest peak during a period of epileptiform activity and that of the standard deviation of a period of low biological activity. Topographic maps of the ECoG signals obtained from the 32 channels were generated by assigning the SNR of each channel to its physical location within the array and normalizing to the channel with the highest SNR for each frame. All data analysis and plotting were performed using a custom MATLAB R2023b code.

**Histological Analyses:** To investigate possible cortical neuronal loss with NeuN (neuronal nuclei) staining and reactive astrogliosis with GFAP (glial fibrillary acidic proteins) levels, rats were sacrificed 3 weeks post-implantation of the ECoG array. Brains were perfused via the transcardiac route with phosphate-buffered saline (PBS) followed by 4% paraformaldehyde in PBS [57, 58].[57, 58]. Brains were then embedded in paraffin, and coronal sections (8 μm) were sampled from the brain region that was in contact with the ECoG array. Histological slides were deparaffinized with xylene and rehydrated in graded alcohol then incubated for 60 minutes in 90 °C sodium citrate buffer for antigen retrieval. Slides were incubated overnight at 4°C with primary antibody followed by peroxidase conjugated secondary antibody for one hour at room temperature (Leica Biosystems, Newcastle, UK). Primary antibodies consisted of anti-NeuN (1:100, Millipore, Burlington, MA, USA) or anti-GFAP (1:250, Santa Cruz Biotechnology, Dallas, TX, USA). Following application of 3,3’-diaminobenzidine (DAB) and hematoxylin counterstaining, images were obtained under light microscopy. NeuN-positive cells were manually counted on six sections per brain to assess cortical neuronal densities (n=2 per group). ImageJ (NIH, Bethesda, MD, USA) was used to measure the optical density for GFAP levels (3 sections per brain, n=2 per group) as well as the cortical surface areas as previously described [32, 59].[32, 59].

### Seizure Classification Model

**Model Development:** Manual annotation of seizure stages was performed by three expert reviewers, assigning one of three class labels to each episode: baseline or low biological activity (Class 0), fast spiking (Class 1), or spike-wave discharges (Class 2). For model development, two representative channels were selected from each recording (E14 and E31) to reduce computational complexity while preserving signal diversity. Channel E14 was characterized by lower SNR, whereas E31 exhibited high SNR, allowing the model to learn from varying signal qualities while avoiding redundancy.

Each signal was segmented using a sliding window of 2 seconds with 50% overlap. After segmentation, each windowed signal was standardized using z-score normalization, as defined in Equation (1):

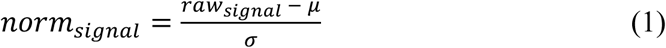

where:

- 𝜇 is the mean of the raw signal segment
- 𝜎 is the standard deviation of the raw signal segment

This normalization ensured consistent scaling across samples (zero mean and unit variance) and improved model convergence for stable model training [60]. The normalized segments and their corresponding labels were saved in “.mat” format for further processing.

**Dataset Partitioning:** In total, 65 minutes of labeled recordings were used for training and testing. The dataset was stratified across two trials, ensuring balance for each class and even distribution between training (70%), validation (10%), and testing (20%) sets. An additional 8-minute recording, unseen during model training, was held out for independent validation. This validation set included all 32 channels and was used to test the final trained models, thereby assessing generalizability across varying signal conditions and electrode placements.

**1D Convolutional Neural Network Architecture:** A one-dimensional (1D) convolutional neural network (CNN) was developed to extract time-domain features from normalized ECoG signals and classify each segment into distinct neural activity stages. Each ECoG segment was treated as a univariate temporal sequence and reshaped into a 4D tensor of shape [N,1,1,samples], where N is the number of time points per window.

The models architecture comprised five convolutional blocks with increasing filter sizes (32, 64, 128, 256, 256). The kernel sizes were initially set to [7, 5, 3] based on standard practice. Each convolutional layer was followed by batch normalization and a rectified linear unit (ReLU) activation. Each filter processes overlapping patches along the temporal axis, enabling the model to extract low-level temporal patterns in early layers, followed by more abstract representations in deeper layers. This formulation is consistent with prior applications of CNNs to EEG data, where input time-series are represented as 1D sequences to preserve their temporal structure during convolution-based feature extraction [61–63]. Max pooling was applied after the first three convolutional layers to progressively reduce the temporal resolution. Dropout regularization (0.3) was introduced after the fourth convolutional block to mitigate overfitting. A global average pooling layer was used to compress the feature space before feeding into the classification head, which consisted of a fully connected layer with 128 ReLU-activated units, followed by dropout (0.5), and a final fully connected layer with softmax activation for three-class prediction.

In addition to the three-class classification problem, binary classification tasks were also investigated to assess discriminability between pairs of classes (0 vs. 1, 0 vs. 2, and 0 vs. 1&2). This multi-stage evaluation allowed us to assess both the overall multiclass performance and the pairwise separability of neural activity states.

**<u>Hyperparameter Optimization:</u>** Hyperparameters were optimized using the training/validation split. To evaluate sensitivity, alternative kernel configurations [3,5,7] and [9,7,5] were tested. In particular, L2 regularization strength with λ values in the range 0.01 to 0.0001 and kernel configurations we systematically varied and tested. The combination yielding the highest validation accuracy was retrained for subsequent testing and reporting. The model was trained using the Adam optimizer for 25 epochs with a mini-batch size of 32. Early stopping was employed during training with termination triggered if validation accuracy failed to improve by at least 0.1% over the previous best value for five consecutive epochs. All training procedures were conducted using MATLAB’s Deep Learning Toolbox (R2024b).

**Model Evaluation:** Performance was first evaluated on the held-out test set using standard metrics, including overall accuracy, macro-averaged F1 score, precision, recall, the average area under the receiver operating characteristic curve (AUC), and confusion matrices.

### Stability and Biodegradability Assessment

The impedance (EIS) of the devices was measured before and after the in vivo surgery to assess the stability of the device during the experiment. The long-term stability of the fabricated devices was studied by continuous monitoring of the impedance of an array immersed in PBS over a period of 3 months at room temperature. For biodegradation assessment, an accelerated chemical hydrolysis in NaOH solution was employed to assess the lifetime of the PCL backbone. The array was fully immersed in an aqueous buffer solution of pH 13 at 37°C.

## References

[1] H. Zhang et al., “Brain–computer interfaces: the innovative key to unlocking neurological conditions,” International Journal of Surgery, vol. 110, no. 9, pp. 5745–5762, 2024, doi: 10.1097/js9.0000000000002022.

[2] J. Wang et al., “Flexible Electrodes for Brain–Computer Interface System,” Advanced Materials, vol. 35, no. 47, p. 2211012, 2023, doi: 10.1002/adma.202211012.

[3] K. J. Miller, D. Hermes, and N. P. StaZ, “The current state of electrocorticography-based brain–computer interfaces,” Neurosurgical focus, vol. 49, no. 1, p. E2, 2020.

[4] A. Kawala-Sterniuk et al., “Summary of over Fifty Years with Brain-Computer Interfaces—A Review,” Brain Sciences, vol. 11, no. 1, p. 43, 2021. [Online]. Available: https://www.mdpi.com/2076-3425/11/1/43.

[5] J. W. Salatino, K. A. Ludwig, T. D. Kozai, and E. K. Purcell, “Glial responses to implanted electrodes in the brain,” Nature biomedical engineering, vol. 1, no. 11, pp. 862–877, 2017.

[6] L. Luan et al., “Recent advances in electrical neural interface engineering: minimal invasiveness, longevity, and scalability,” Neuron, vol. 108, no. 2, pp. 302–321, 2020.

[7] B. Zhang, C. Deng, C. Cai, and X. Li, “In vivo neural interfaces—from small-to large-scale recording,” Frontiers in Nanotechnology, vol. 4, p. 885411, 2022.

[8] M. E. E. Alahi, Y. Liu, Z. Xu, H. Wang, T. Wu, and S. C. Mukhopadhyay, “Recent advancement of electrocorticography (ECoG) electrodes for chronic neural recording/stimulation,” Materials Today Communications, vol. 29, p. 102853, 2021.

[9] J. L. Roland, C. D. Hacker, and E. C. Leuthardt, “A review of passive brain mapping techniques in neurological surgery,” Neurosurgery, vol. 88, no. 1, pp. 15–24, 2021.

[10] U.-J. Jeong et al., “A minimally invasive flexible electrode array for simultaneous recording of ECoG signals from multiple brain regions,” Lab on a Chip, vol. 21, no. 12, pp. 2383–2397, 2021.

[11] F. R. Jahangiri, A. Dobariya, A. Kruse, O. Kalyta, and J. D. Moorman, “Mapping of the motor cortex,” Cureus, vol. 12, no. 9, 2020.

[12] Y. Xie et al., “Materials and devices for high-density, high-throughput micro-electrocorticography arrays,” Fundamental Research, vol. 5, no. 1, pp. 17–28, 2025/01/01/ 2025, doi: 10.1016/j.fmre.2024.01.016.

[13] Y. Chen, Y. Zhang, Z. Liang, Y. Cao, Z. Han, and X. Feng, “Flexible inorganic bioelectronics,” npj Flexible Electronics, vol. 4, no. 1, p. 2, 2020/02/04 2020, doi: 10.1038/s41528-020-0065-1.

[14] J. Parvizi and S. Kastner, “Promises and limitations of human intracranial electroencephalography,” Nature Neuroscience, vol. 21, no. 4, pp. 474–483, 2018/04/01 2018, doi: 10.1038/s41593-018-0108-2.

[15] X. Liu, Y. Gong, Z. Jiang, T. Stevens, and W. Li, “Flexible high-density microelectrode arrays for closed-loop brain–machine interfaces: a review,” Frontiers in Neuroscience, vol. 18, p. 1348434, 2024.

[16] X. Wang, A. Aziz, X. Sheng, L. Wang, and L. Yin, “Bioresorbable neural interfaces for bioelectronic medicine,” Current Opinion in Biomedical Engineering, vol. 32, p. 100565, 2024/12/01/ 2024, doi: 10.1016/j.cobme.2024.100565.

[17] S. Wei et al., “Shape-changing electrode array for minimally invasive large-scale intracranial brain activity mapping,” Nature Communications, vol. 15, no. 1, p. 715, 2024/01/24 2024, doi: 10.1038/s41467-024-44805-2.

[18] F. Au - Fallegger, A. Au - Trouillet, and S. P. Au - Lacour, *JoVE*, no. 193, p. e64997, 2023/03/31/ 2023, doi: doi:10.3791/64997.

[19] R. Matta, D. Moreau, and R. O’Connor, “Printable devices for neurotechnology,” Frontiers in Neuroscience, vol. 18, p. 1332827, 2024.

[20] G. Baldini, A. Albini, P. Maiolino, and G. Cannata, “An Atlas for the Inkjet Printing of Large-Area Tactile Sensors,” Sensors, vol. 22, no. 6, p. 2332, 2022. [Online]. Available: https://www.mdpi.com/1424-8220/22/6/2332.

[21] A. Imai et al., “Flexible Thin-Film Neural Electrodes with Improved Conformability for ECoG Measurements and Electrical Stimulation,” Advanced Materials Technologies, vol. 8, no. 21, p. 2300300, 2023, doi: 10.1002/admt.202300300.

[22] S. Oribe et al., “Hydrogel-based organic subdural electrode with high conformability to brain surface,” Scientific Reports, vol. 9, no. 1, p. 13379, 2019.

[23] P. D. Donaldson, L. Ghanbari, M. L. Rynes, S. B. Kodandaramaiah, and S. L. Swisher, “Inkjet-printed silver electrode array for in-vivo electrocorticography,” in 2019 9th International IEEE/EMBS Conference on Neural Engineering (NER), 2019: IEEE, pp. 774–777.

[24] L. Liu et al., “Implantable Brain-Computer Interface Based On Printing Technology,” in 2023 11th International Winter Conference on Brain-Computer Interface (BCI), 20-22 Feb. 2023 2023, pp. 1–5, doi: 10.1109/BCI57258.2023.10078643.

[25] Y. Kim, S. Alimperti, P. Choi, and M. Noh, “An inkjet printed flexible Electrocorticography (ECoG) microelectrode Array on a thin Parylene-C film,” Sensors, vol. 22, no. 3, p. 1277, 2022.

[26] Y. Xie et al., “Materials and devices for high-density, high-throughput micro-electrocorticography arrays,” Fundamental Research, 2024/02/28/ 2024, doi: 10.1016/j.fmre.2024.01.016.

[27] Z. Aqrawe et al., “A simultaneous optical and electrical in-vitro neuronal recording system to evaluate microelectrode performance,” PLOS ONE, vol. 15, no. 8, p. e0237709, 2020, doi: 10.1371/journal.pone.0237709.

[28] D. W. Park et al., “Graphene-based carbon-layered electrode array technology for neural imaging and optogenetic applications,” (in eng), Nat Commun, vol. 5, p. 5258, Oct 20 2014, doi: 10.1038/ncomms6258.

[29] M. E. E. Alahi et al., “Slippery Epidural ECoG Electrode for High-Performance Neural Recording and Interface,” Biosensors, vol. 12, no. 11, p. 1044, 2022. [Online]. Available: https://www.mdpi.com/2079-6374/12/11/1044.

[30] Y. Nam and B. C. Wheeler, “In vitro microelectrode array technology and neural recordings,” Critical Reviews™ in Biomedical Engineering, vol. 39, no. 1, 2011.

[31] F. Khoury, S. Saleh, H. Badawe, M. Obeid, and M. Khraiche, “Inkjet-printed, flexible organic electrochemical transistors for high-performance electrocorticography recordings,” ACS Applied Materials & Interfaces, vol. 16, no. 41, pp. 55045–55055, 2024.

[32] Y. Mrad et al., “Lestaurtinib (CEP-701) reduces the duration of limbic status epilepticus in periadolescent rats,” Epilepsy Research, vol. 195, p. 107198, 2023.

[33] R. J. Staba and A. Bragin, “High-frequency oscillations and other electrophysiological biomarkers of epilepsy: underlying mechanisms,” Biomarkers in medicine, vol. 5, no. 5, pp. 545–556, 2011.

[34] J. Hu et al., “Fully desktop fabricated flexible graphene electrocorticography (ECoG) arrays,” Journal of neural engineering, vol. 20, no. 1, p. 016019, 2023.

[35] R. Dong et al., “Printed stretchable liquid metal electrode arrays for in vivo neural recording,” Small, vol. 17, no. 14, p. 2006612, 2021.

[36] R. M. Almasri, W. AlChamaa, A. R. Tehrani-Bagha, and M. L. Khraiche, “Highly Flexible Single-Unit Resolution All Printed Neural Interface on a Bioresorbable Backbone,” ACS Applied Bio Materials, vol. 3, no. 10, pp. 7040–7051, 2020/10/19 2020, doi: 10.1021/acsabm.0c00895.

[37] H. K. Meeren, J. P. M. Pijn, E. L. Van Luijtelaar, A. M. Coenen, and F. H. L. da Silva, “Cortical focus drives widespread corticothalamic networks during spontaneous absence seizures in rats,” Journal of Neuroscience, vol. 22, no. 4, pp. 1480–1495, 2002.

[38] M. Avoli, M. de Curtis, and R. Köhling, “Does interictal synchronization influence ictogenesis?,” Neuropharmacology, vol. 69, pp. 37–44, 2013.

[39] K. Xu et al., “Bioresorbable Electrode Array for Electrophysiological and Pressure Signal Recording in the Brain,” (in eng), Adv Healthc Mater, vol. 8, no. 15, p. e1801649, Aug 2019, doi: 10.1002/adhm.201801649.

[40] H. Moon, J.-W. Jang, S. Park, J.-H. Kim, J. S. Kim, and S. Kim, “Soft, conformal PDMS-based ECoG electrode array for long-term in vivo applications,” Sensors and Actuators B: Chemical, vol. 401, p. 135099, 2024.

[41] Y. Liu et al., “Flexible, high-density, laminated ECoG electrode array for high spatiotemporal resolution foci diagnostic localization of refractory epilepsy,” Bio-Design and Manufacturing, vol. 7, no. 4, pp. 388–398, 2024.

[42] M. Edoho, C. Mooney, and L. Wei, “AI-Based Electroencephalogram Analysis in Rodent Models of Epilepsy: A Systematic Review,” Applied Sciences, vol. 14, no. 16, p. 7398, 2024.

[43] W. Barry, S. Arcot Desai, T. K. Tcheng, and M. J. Morrell, “A high accuracy electrographic seizure classifier trained using semi-supervised labeling applied to a large spectrogram dataset,” Frontiers in neuroscience, vol. 15, p. 667373, 2021.

[44] L. Kamintsky et al., “An algorithm for seizure detection in rodents,” Epilepsia Open, 2025.

[45] M. Hartmann et al., “Seizure detection with deep neural networks for review of two-channel electroencephalogram,” Epilepsia, vol. 64, pp. S34–S39, 2023.

[46] M. Wu et al., “Ultrathin, soft, bioresorbable organic electrochemical transistors for transient spatiotemporal mapping of brain activity,” Advanced Science, vol. 10, no. 14, p. 2300504, 2023.

[47] J.-y. Liou et al., “A model for focal seizure onset, propagation, evolution, and progression,” Elife, vol. 9, p. e50927, 2020.

[48] J. Viventi et al., “Flexible, foldable, actively multiplexed, high-density electrode array for mapping brain activity in vivo,” Nature neuroscience, vol. 14, no. 12, pp. 1599–1605, 2011.

[49] L. F. Rossi, R. C. Wykes, D. M. Kullmann, and M. Carandini, “Focal cortical seizures start as standing waves and propagate respecting homotopic connectivity,” Nature communications, vol. 8, no. 1, p. 217, 2017.

[50] N. Natraj et al., “Sampling representational plasticity of simple imagined movements across days enables long-term neuroprosthetic control,” Cell, vol. 188, no. 5, pp. 1208–1225.e32, 2025, doi: 10.1016/j.cell.2025.02.001.

[51] N. Natraj, et al., “Flexible regulation of representations on a drifting manifold enables long-term stable complex neuroprosthetic control,” (in eng), bioRxiv, Aug 14 2023, doi: 10.1101/2023.08.11.551770.

[52] H.-S. Kim, S. R. Dhage, D.-E. Shim, and H. T. Hahn, “Intense pulsed light sintering of copper nanoink for printed electronics,” Applied Physics A, vol. 97, pp. 791–798, 2009.

[53] T. Falat, J. Felba, B. Platek, T. Piasecki, A. Moscicki, and A. Smolarek, “Low-temperature, photonic approach to sintering the ink-jet printed conductive microstructures containing nano sized silver particles,” in 18th European Microelectronics & Packaging Conference, 2011: IEEE, pp. 1–4.

[54] M. Trotter et al., “Inkjet-Printing of Nanoparticle Gold and Silver Ink on Cyclic Olefin Copolymer for DNA-Sensing Applications,” (in eng), Sensors (Basel), vol. 20, no. 5, Feb 29 2020, doi: 10.3390/s20051333.

[55] C. E. Hajjaji, N. Delhote, S. Verdeyme, M. Piechowiak, and O. Durand, “Optimization of the conductivity of microwave components printed by inkjet on polymeric substrates by photonic sintering,” in 2020 50th European Microwave Conference (EuMC), 12-14 Jan. 2021 2021, pp. 380–383, doi: 10.23919/EuMC48046.2021.9338007.

[56] A. Sharif, N. Farid, and G. M. O’Connor, “Ultrashort laser sintering of metal nanoparticles: A review,” Results in Engineering, vol. 16, p. 100731, 2022/12/01/ 2022, doi: 10.1016/j.rineng.2022.100731.

[57] R. Asdikian et al., “Hippocampal injury and learning deficits following non-convulsive status epilepticus in periadolescent rats,” Epilepsy & Behavior, vol. 125, p. 108415, 2021.

[58] D. Jalloul et al., “Potentiating hemorrhage in a periadolescent rat model of closed-head traumatic brain injury worsens hyperexcitability but not behavioral deficits,” Int J Mol Sci, vol. 22, no. 12, p. 6456, 2021.

[59] Y. Medlej et al., “Lestaurtinib (CEP-701) modulates the eZects of early life hypoxic seizures on cognitive and emotional behaviors in immature rats,” Epilepsy & Behavior, vol. 92, pp. 332–340, 2019.

[60] D. Thara and B. PremaSudha, “Auto-detection of epileptic seizure events using deep neural network with diZerent feature scaling techniques,” Pattern Recognition Letters, vol. 128, pp. 544–550, 2019.

[61] I. Rakhmatulin, M.-S. Dao, A. Nassibi, and D. Mandic, “Exploring convolutional neural network architectures for EEG feature extraction,” Sensors, vol. 24, no. 3, p. 877, 2024.

[62] G. Xu, T. Ren, Y. Chen, and W. Che, “A one-dimensional CNN-LSTM model for epileptic seizure recognition using EEG signal analysis,” Frontiers in neuroscience, vol. 14, p. 578126, 2020.

[63] X. Wang, X. Wang, W. Liu, Z. Chang, T. Kärkkäinen, and F. Cong, “One dimensional convolutional neural networks for seizure onset detection using long-term scalp and intracranial EEG,” Neurocomputing, vol. 459, pp. 212–222, 2021.

