## Supplementary for "“Rapid prototyping of flexible biodegradable ECoG arrays for high-resolution cortical mapping and real-time seizure classification ”"

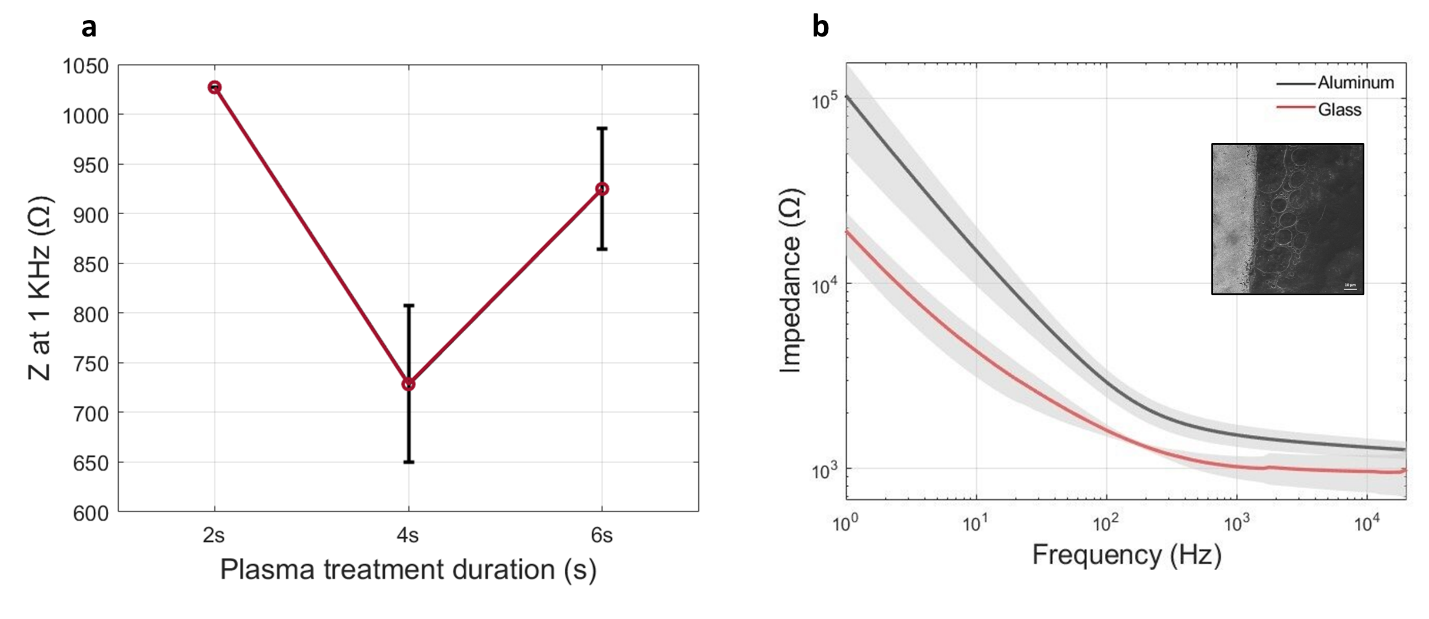

Supplementary Figure 1. (a) Effect of plasma treatment duration on electrode impedance magnitude at 1 kHz. (b) Averaged impedance spectrograms of electrodes printed on PCL with a glass vs aluminum backbone. Inset: SEM image of gold nanoparticles on PCL film with aluminum backbone showing damage to PCL film.

Supplementary Table 1. Key developments of flexible microelectrode array platforms for electrophysiological monitoring

| Author | Year | Patterning method | Electrode material | Substrate | Electrode size (um) | Electrode number | Sensing area (mm^2^) | Electrode Density (electrode/mm^2^) | Impedance (KΩ at 1 KHz) | Thickness (µm) | In-vivo testing | SNR (dB) |
| --- | --- | --- | --- | --- | --- | --- | --- | --- | --- | --- | --- | --- |
| This work | 2025 | Inkjet printing | Au nanoparticles | PCL film | 75 | 36 | 4.84 | 7.44 | 10.6 | ~10 | rat | 28 |
| Moon [1] | 2024 | Photolithography | Ti/Au | Parylene-filled PDMS | 150 | 16 | 7.5 | 2.13 | 27.8 | 80 | rat & monkey | - |
| Liu [2] | 2024 | Photolithography | Cr/Au | Polyimide | <15 | 800 | 0.18 | 4444 | 5.3 | 30 | rabbit | - |
| Hu [3] | 2023 | Stencil printing | Graphene | Polyimide | 300 | 16 | 36 | 0.44 | 24.72 | - | rat | - |
| Imai [4] | 2023 | Inkjet printing | Au nanoparticles | SBS | 100 | 35 | 56* | 0.625 | 1.68 | 8 | rat | 18.2 |
| Kim [5] | 2022 | Inkjet printing | AgNP | Parylene C | 100 | 20 | 4.75* | 4.21 | 90.23 | 10 | rat & human | - |
| Li [6] | 2021 | Photolithography | Au-MWCNTs/PEDOT:PSS | PDMS/Parylene C | 60 | 14 | 9.73 | 1.71 | 20.68 | 20 | rat | 10.05* |
| Dong [7] | 2021 | Screen printing | Eutectic gallium–indium (EGaIn) | PDMS | 100 | 16 | 9.73 | 1.64 | 250 | 100 | rat | - |
| Kaiju [8] | 2021 | Photolithography | Pt black | Parylene C | 50 | 1152 | 98 | 11.8 | 26 | 20 | monkey | - |
| Patil [9] | 2020 | Photolithography | Au | Silk | 250 | 12 | 2.25 | 5.33 | 20.57 | ~30 | rat | - |
| Seo [10] | 2020 | Photolithography | Au Nanonetwork | Polyimide | 200 | 16 | 6.76 | 2.37 | 9.1 | 5 | mouse | ~11* |
| Borda [11] | 2020 | Inkjet printing | Platinum | PI | 450 | 16 | ~14.8* | 1.08 | 9.6 | 60 | rabbit | - |
| Donaldson [12] | 2019 | Inkjet printing | AgNP | PET | 300 | 6 | 4 | 1.5 | ~1.5* | 50 | mouse | ~37 |
| Xu [13] | 2019 | Magnetron sputtering/mask | Mo | PLLA/PCL | 200 | 6 | 2.2* | 2.73 | 10.2 | 40 | rat | ~4* |
| Shi [14] | 2019 | Photolithography | Au | PI/Silk | 250 | 100 | 100 | 0.49 | - | 30 | rat | <4* |
| Ganji [15] | 2019 | Photolithography | Pt Nanorods | Parylene C | 50 | 128 | 0.23 | 568 | 16.89 | - | songbird, monkey & mouse | - |
| Schander [16] | 2019 | Photolithography | Au/PEDOT:PSS | Polyimide | 560 | 202 | 780 | 0.26 | 1.1 | 10 | monkey | - |
| Garma | 2019 | Inkjet printing | PEDOT:PSS | Polyimide | 300 | 16 | - | - | 19.5 | 25 | - | 10.2 |
| Oribe [17] | 2019 | Electropolymerization | PEDOT-CF | PVA hydrogel | 3000 | - | - | - | ~0.02* | 1000 | rat | 10 |
| Vitale [18] | 2018 | Photolithography | Au/ECM | Parylene C/ECM | 50 | 8 | 45 | 0.18 | 190 | 30 | rat | - |
| Guo [19] | 2018 | Photolithography | Au | Nanopaper | 150 | 9 | ~2* | 4.5 | 200 | 40 | rat | - |
| Zhang [20] | 2018 | Photolithography | CNT | PDMS | 100 | 16 | 9* | 1.78 | 200 | 100 | rat | - |
| Tybrandt [21] | 2018 | Photolithography | Au-TiO2 Nanowires | PDMS | 50 | 32 | ~0.84 | 38 | 10 | ~80 | rat | - |
| Yu [22] | 2016 | Photolithography | SiO2−Mo | PLGA | 250 | 4 | 12 | 0.33 | ~20* | 30 | rat | - |
| Khodagholy [23] | 2016 | Photolithography | PEDOT:PSS | Parylene C | 10 | 240 | 840 | 0.29 | ~30 | 4 | human | - |
| Khodagholy [24] | 2015 | Photolithography | PEDOT:PSS | Parylene C | 10 | 64 | 0.04 | 1600 | 30 | 4 | rat & human | - |
| Park [25] | 2014 | Photolithography | Graphene | Parlyene C | 200 | 16 | 3.61 | 4.43 | 243.5 | - | rat & mouse | - |
| Khodagholy [26] | 2013 | Photolithography | Au/PEDOT:PSS | Parylene C | 12 | 8 | - | - | - | 4 | rat | 24.2 |
| Ledochowitsch [27] | 2013 | Photolithography | Pt black | Parlyene C | 40 | 64 | 1.96 | 32.7 | - |  | rat | - |
| Toda [28] | 2011 | Photolithography | Pt Black | Parlyene C | 50 | 32 | 36 | 0.89 | 103 | 20 | rat | - |
| Ledochowitsch [29] | 2011 | Photolithography | Pt | Parylene C | 440 | 256 | 56 | 4.6 | ~11 | 12 | rat | - |
| Kim [30] | 2010 | Photolithography | Au | PI/Silk | 500 | 30 | 80* | 0.38 | - | - | cat | - |

Supplementary Table 2: Performance Metrics Across All Models

| Model | Accuracy | Macro-F1 | Macro-Precision | Macro-Recall | Average AUC |
| --- | --- | --- | --- | --- | --- |
| Three-Class | 93.1% | 93.1% | 93.8% | 93.1% | 98.7% |
| Combined | 97.7% | 97.4% | 97.0% | 97.9% | 99.4% |
| Binary 0 vs 1 | 95.8% | 95.8% | 96.0% | 95.8% | 99.9% |
| Binary 0 vs 2 | 99.6% | 99.6% | 99.6% | 99.6% | 100.0% |

Supplementary Table 3: Confusion Matrix of the Three-Class Model

| True Class | Baseline | 123 | 5 | 2 |
| --- | --- | --- | --- | --- |
|  | Fast spiking | 1 | 128 | 1 |
|  | Spike-wave discharge | 0 | 18 | 112 |
|  |  | Baseline | Fast spiking | Spike-wave discharge |
|  |  | Predicted Class | | |

Supplementary Table 4: Validation of the Three-Class Model on 8-Minute Recording Across All Channels

| Channel | Accuracy | Macro-F1 | Macro-Precision | Macro-Recall | Average AUC |
| --- | --- | --- | --- | --- | --- |
| E1 | 90.8% | 90.9% | 91.3% | 90.8% | 99.2% |
| E2 | 90.8% | 90.9% | 91.8% | 90.8% | 99.2% |
| E3 | 90.4% | 90.5% | 91.0% | 90.4% | 99.0% |
| E4 | 85.8% | 85.5% | 86.6% | 85.8% | 97.9% |
| E5 | 93.8% | 93.8% | 94.0% | 93.8% | 99.2% |
| E6 | 93.3% | 93.3% | 93.4% | 93.3% | 98.8% |
| E7 | 95.0% | 95.0% | 95.2% | 95.0% | 99.2% |
| E8 | 90.8% | 90.8% | 91.5% | 90.8% | 98.9% |
| E9 | 93.8% | 93.8% | 93.9% | 93.8% | 99.2% |
| E10 | 91.3% | 91.2% | 91.5% | 91.3% | 98.6% |
| E11 | 94.6% | 94.6% | 94.6% | 94.6% | 99.4% |
| E12 | 95.8% | 95.8% | 96.0% | 95.8% | 99.3% |
| E13 | 41.3% | 29.7% | 29.0% | 41.3% | 60.6% |
| E14 | 95.8% | 95.8% | 95.9% | 95.8% | 99.3% |
| E15 | 37.1% | 23.9% | 44.9% | 37.1% | 72.1% |
| E16 | 83.3% | 83.1% | 87.8% | 83.3% | 96.8% |
| E17 | 73.8% | 72.4% | 84.3% | 73.8% | 94.4% |
| E18 | 79.2% | 78.9% | 85.9% | 79.2% | 96.6% |
| E19 | 85.4% | 85.6% | 89.0% | 85.4% | 97.2% |
| E20 | 82.9% | 83.0% | 87.8% | 82.9% | 96.9% |
| E21 | 35.8% | 21.6% | 44.7% | 35.8% | 68.5% |
| E22 | 88.3% | 88.5% | 90.4% | 88.3% | 97.6% |
| E23 | 90.0% | 90.1% | 92.1% | 90.0% | 97.5% |
| E24 | 92.9% | 93.0% | 94.0% | 92.9% | 98.6% |
| E25 | 93.8% | 93.7% | 93.8% | 93.8% | 98.6% |
| E26 | 92.9% | 92.9% | 93.5% | 92.9% | 98.4% |
| E27 | 92.5% | 92.5% | 93.0% | 92.5% | 98.3% |
| E28 | 93.3% | 93.3% | 93.3% | 93.3% | 98.4% |
| E29 | 93.3% | 93.3% | 93.4% | 93.3% | 98.8% |
| E30 | 94.2% | 94.2% | 94.2% | 94.2% | 98.7% |
| E31 | 93.3% | 93.3% | 93.3% | 93.3% | 98.4% |
| E32 | 93.3% | 93.3% | 93.3% | 93.3% | 98.3% |
| Average | 85.6% | 84.3% | 87.0% | 85.6% | 95.4% |

Supplementary Table 5: Validation of the Combined Model on 8-Minute Recording Across All Channels

| Channel | Accuracy | Macro-F1 | Macro-Precision | Macro-Recall | Average AUC |
| --- | --- | --- | --- | --- | --- |
| U1 | 89.6% | 88.8% | 87.7% | 90.6% | 97.1% |
| U2 | 90.4% | 89.7% | 88.6% | 91.6% | 97.2% |
| U3 | 88.8% | 88.0% | 87.0% | 90.3% | 97.7% |
| U4 | 70.0% | 69.9% | 75.8% | 77.2% | 95.9% |
| U5 | 97.9% | 97.7% | 97.3% | 98.1% | 99.9% |
| U6 | 78.8% | 78.3% | 79.5% | 83.1% | 97.4% |
| U7 | 87.5% | 86.9% | 86.2% | 90.3% | 99.2% |
| U8 | 69.6% | 69.5% | 75.1% | 76.6% | 97.0% |
| U9 | 81.3% | 80.7% | 81.1% | 85.0% | 97.6% |
| U10 | 76.3% | 75.9% | 78.0% | 81.3% | 96.6% |
| U11 | 79.1% | 78.7% | 80.4% | 84.0% | 97.7% |
| U12 | 81.3% | 80.8% | 81.7% | 85.6% | 98.4% |
| U13 | 50.4% | 50.4% | 55.4% | 55.6% | 57.7% |
| U14 | 80.8% | 80.4% | 81.4% | 85.3% | 98.3% |
| U15 | 79.2% | 72.2% | 82.5% | 70.3% | 86.0% |
| U16 | 94.6% | 93.9% | 93.8% | 94.1% | 98.6% |
| U17 | 96.3% | 95.7% | 97.3% | 94.4% | 98.7% |
| U18 | 95.0% | 94.3% | 94.6% | 94.1% | 98.6% |
| U19 | 95.0% | 94.3% | 94.9% | 93.8% | 98.0% |
| U20 | 94.6% | 93.8% | 94.3% | 93.4% | 98.2% |
| U21 | 73.8% | 61.1% | 79.7% | 61.6% | 73.1% |
| U22 | 94.6% | 93.8% | 94.6% | 93.1% | 98.0% |
| U23 | 95.4% | 94.9% | 94.5% | 95.3% | 98.5% |
| U24 | 95.8% | 95.3% | 95.1% | 95.6% | 98.4% |
| U25 | 81.7% | 80.9% | 80.7% | 84.4% | 94.7% |
| U26 | 95.8% | 95.3% | 95.9% | 94.7% | 98.6% |
| U27 | 96.7% | 96.3% | 96.3% | 96.3% | 98.6% |
| U28 | 80.8% | 80.1% | 80.1% | 83.8% | 93.9% |
| U29 | 89.6% | 88.7% | 87.7% | 90.3% | 95.2% |
| U30 | 89.2% | 88.2% | 87.3% | 89.7% | 94.9% |
| U31 | 80.0% | 79.2% | 79.2% | 82.8% | 93.5% |
| U32 | 79.6% | 78.8% | 78.9% | 82.5% | 93.5% |
| Average | 85.3% | 84.1% | 85.7% | 86.4% | 94.9% |

References

[1]	H. Moon, J.-W. Jang, S. Park, J.-H. Kim, J. S. Kim, and S. Kim, "Soft, conformal PDMS-based ECoG electrode array for long-term in vivo applications," *Sensors and Actuators B: Chemical,* vol. 401, p. 135099, 2024.

[2]	Y. Liu *et al.*, "Flexible, high-density, laminated ECoG electrode array for high spatiotemporal resolution foci diagnostic localization of refractory epilepsy," *Bio-Design and Manufacturing,* vol. 7, no. 4, pp. 388-398, 2024.

[3]	J. Hu *et al.*, "Fully desktop fabricated flexible graphene electrocorticography (ECoG) arrays," *Journal of neural engineering,* vol. 20, no. 1, p. 016019, 2023.

[4]	A. Imai *et al.*, "Flexible Thin-Film Neural Electrodes with Improved Conformability for ECoG Measurements and Electrical Stimulation," *Advanced Materials Technologies,* vol. 8, no. 21, p. 2300300, 2023, doi: <https://doi.org/10.1002/admt.202300300>.

[5]	Y. Kim, S. Alimperti, P. Choi, and M. Noh, "An inkjet printed flexible Electrocorticography (ECoG) microelectrode Array on a thin Parylene-C film," *Sensors,* vol. 22, no. 3, p. 1277, 2022.

[6]	X. Li *et al.*, "PDMS–parylene hybrid, flexible micro-ECoG electrode array for spatiotemporal mapping of epileptic electrophysiological activity from multicortical brain regions," *ACS Applied Bio Materials,* vol. 4, no. 11, pp. 8013-8022, 2021.

[7]	R. Dong *et al.*, "Printed stretchable liquid metal electrode arrays for in vivo neural recording," *Small,* vol. 17, no. 14, p. 2006612, 2021.

[8]	T. Kaiju, M. Inoue, M. Hirata, and T. Suzuki, "High-density mapping of primate digit representations with a 1152-channel µECoG array," *Journal of Neural Engineering,* vol. 18, no. 3, p. 036025, 2021.

[9]	A. C. Patil, A. Bandla, Y.-H. Liu, B. Luo, and N. V. Thakor, "Nontransient silk sandwich for soft, conformal bionic links," *Materials Today,* vol. 32, pp. 68-83, 2020.

[10]	J. W. Seo *et al.*, "Artifact‐free 2D mapping of neural activity in vivo through transparent gold nanonetwork array," *Advanced Functional Materials,* vol. 30, no. 34, p. 2000896, 2020.

[11]	E. Borda *et al.*, "All‐printed electrocorticography array for in vivo neural recordings," *Advanced Engineering Materials,* vol. 22, no. 3, p. 1901403, 2020.

[12]	P. D. Donaldson, L. Ghanbari, M. L. Rynes, S. B. Kodandaramaiah, and S. L. Swisher, "Inkjet-printed silver electrode array for in-vivo electrocorticography," in *2019 9th International IEEE/EMBS Conference on Neural Engineering (NER)*, 2019: IEEE, pp. 774-777.

[13]	K. Xu *et al.*, "Bioresorbable Electrode Array for Electrophysiological and Pressure Signal Recording in the Brain," (in eng), *Adv Healthc Mater,* vol. 8, no. 15, p. e1801649, Aug 2019, doi: 10.1002/adhm.201801649.

[14]	Z. Shi *et al.*, "Silk‐enabled conformal multifunctional bioelectronics for investigation of spatiotemporal epileptiform activities and multimodal neural encoding/decoding," *Advanced Science,* vol. 6, no. 9, p. 1801617, 2019.

[15]	M. Ganji *et al.*, "Selective formation of porous Pt nanorods for highly electrochemically efficient neural electrode interfaces," *Nano letters,* vol. 19, no. 9, pp. 6244-6254, 2019.

[16]	A. Schander, S. Strokov, H. Stemmann, T. Teßmann, A. K. Kreiter, and W. Lang, "A flexible 202-channel epidural ECoG array with PEDOT: PSS coated electrodes for chronic recording of the visual cortex," *IEEE Sensors Journal,* vol. 19, no. 3, pp. 820-825, 2018.

[17]	S. Oribe *et al.*, "Hydrogel-based organic subdural electrode with high conformability to brain surface," *Scientific Reports,* vol. 9, no. 1, p. 13379, 2019.

[18]	F. Vitale *et al.*, "Biomimetic extracellular matrix coatings improve the chronic biocompatibility of microfabricated subdural microelectrode arrays," *PLoS One,* vol. 13, no. 11, p. e0206137, 2018.

[19]	Y. Guo *et al.*, "Flexible and biocompatible nanopaper-based electrode arrays for neural activity recording," *Nano Research,* vol. 11, pp. 5604-5614, 2018.

[20]	J. Zhang *et al.*, "Stretchable transparent electrode arrays for simultaneous electrical and optical interrogation of neural circuits in vivo," *Nano letters,* vol. 18, no. 5, pp. 2903-2911, 2018.

[21]	K. Tybrandt *et al.*, "High‐density stretchable electrode grids for chronic neural recording," *Advanced Materials,* vol. 30, no. 15, p. 1706520, 2018.

[22]	K. J. Yu *et al.*, "Bioresorbable silicon electronics for transient spatiotemporal mapping of electrical activity from the cerebral cortex," *Nature materials,* vol. 15, no. 7, pp. 782-791, 2016.

[23]	D. Khodagholy *et al.*, "Organic electronics for high-resolution electrocorticography of the human brain," *Science Advances,* vol. 2, no. 11, p. e1601027, 2016.

[24]	D. Khodagholy *et al.*, "NeuroGrid: recording action potentials from the surface of the brain," *Nature neuroscience,* vol. 18, no. 2, pp. 310-315, 2015.

[25]	D. W. Park *et al.*, "Graphene-based carbon-layered electrode array technology for neural imaging and optogenetic applications," (in eng), *Nat Commun,* vol. 5, p. 5258, Oct 20 2014, doi: 10.1038/ncomms6258.

[26]	D. Khodagholy *et al.*, "In vivo recordings of brain activity using organic transistors," *Nature communications,* vol. 4, no. 1, p. 1575, 2013.

[27]	P. Ledochowitsch, A. C. Koralek, D. Moses, J. M. Carmena, and M. M. Maharbiz, "Sub-mm functional decoupling of electrocortical signals through closed-loop BMI learning," in *2013 35th Annual International Conference of the IEEE Engineering in Medicine and Biology Society (EMBC)*, 2013: IEEE, pp. 5622-5625.

[28]	H. Toda, T. Suzuki, H. Sawahata, K. Majima, Y. Kamitani, and I. Hasegawa, "Simultaneous recording of ECoG and intracortical neuronal activity using a flexible multichannel electrode-mesh in visual cortex," *Neuroimage,* vol. 54, no. 1, pp. 203-212, 2011.

[29]	P. Ledochowitsch, R. Félus, R. Gibboni, A. Miyakawa, S. Bao, and M. Maharbiz, "Fabrication and testing of a large area, high density, parylene MEMS µECoG array," in *2011 IEEE 24th International Conference on Micro Electro Mechanical Systems*, 2011: IEEE, pp. 1031-1034.

[30]	D.-H. Kim *et al.*, "Dissolvable films of silk fibroin for ultrathin conformal bio-integrated electronics," *Nature materials,* vol. 9, no. 6, pp. 511-517, 2010.
